# Diet suppresses tumour initiation by maintaining quiescence of mutation-bearing neural stem cells

**DOI:** 10.1101/2022.03.09.483611

**Authors:** Valeria Amodeo, Timothy Davies, Amalia Martinez-Segura, Melanie Clements, Holly Simpson Ragdale, Andrew Bailey, Mariana Silva Dos Santos, James I. MacRae, Joao Mokochinski, Holger Kramer, Claudia Garcia-Diaz, Alex P. Gould, Samuel Marguerat, Simona Parrinello

**Author notes:** these two authors contributed equally to the work (Timothy Davies, Amalia Martinez-Segura).

## Abstract

Glioblastoma is thought to originate from neural stem cells (NSCs) of the subventricular zone that acquire genetic alterations. In the adult brain, NSCs are largely quiescent, suggesting that deregulation of quiescence maintenance may be a pre-requisite for tumour initiation. Although inactivation of the tumour suppressor p53 is a frequent event in gliomagenesis, whether, or how, it affects quiescent NSCs (qNSCs) remains unclear. Here we show that p53 maintains quiescence by inducing fatty acid oxidation (FAO) and that acute p53 deletion in qNSCs results in their premature activation to a proliferative state. Mechanistically, this occurs through direct transcriptional induction of PPARGC1a, which in turn activates PPARα to upregulate FAO genes. Strikingly, dietary supplementation with fish oil containing omega-3 fatty acids, natural PPARα ligands, fully restores quiescence of p53-deficient NSCs and delays tumour initiation in a glioblastoma mouse model. Thus, diet can silence glioblastoma driver mutations, with important implications for cancer prevention.

## Introduction

Increasing evidence indicates that glioblastoma (GBM) originates from NSCs of the subventricular zone (SVZ), one of the two main neurogenic niches of the adult brain (Sanai *et al*., 2004; Lee *et al*., 2018). SVZ NSCs, which are largely quiescent (qNSCs) in the normal adult brain (Sanai *et al*., 2004), were shown to bear mutations in cancer-driving genes such as *TERT, TP53* and *EGFR* in GBM patients (Lee *et al*., 2018). This suggests that these genes may play a role in controlling transitions between quiescence and activation to a proliferative state at the onset of tumourigenesis. Therefore, deciphering whether and how GBM-relevant mutations affect the biology of qNSCs should provide fundamental understanding of disease aetiology and reveal novel approaches for cancer prevention.

The transcription factor p53 (*TP53* in humans, *Trp53* in mice) is the most frequently mutated gene in human cancer and the p53 pathway is altered in 87% of GBM patients (Brennan *et al*., 2013). p53 exerts its tumour suppressive effects through the regulation of a myriad of transcriptional programmes, including canonical suppression of apoptosis, cell-cycle and senescence, as well as non-canonical modulation of metabolism, pluripotency and autophagy (Bieging, Mello and Attardi, 2014; Labuschagne, Zani and Vousden, 2018). Interestingly, p53 has also been linked to the regulation of murine adult SVZ neurogenesis (Gil-Perotin *et al*., 2006; Meletis *et al*., 2006). In rodents, neurogenesis occurs from a subpopulation of NSCs termed type-B cells (Lim and Alvarez-Buylla, 2016). Although largely quiescent, adult murine type-B cells retain ability to activate to a proliferative state (aNSCs) from which they give rise to transit amplifying progenitors (type-C cells) that in turn fuel the production of neurons and glia (Lim and Alvarez-Buylla, 2016). Analysis of constitutive p53 knock-out mouse models has shown that p53 restrains type-C cell proliferation and neuronal differentiation (Gil-Perotin *et al*., 2006; Meletis *et al*., 2006). In contrast, its role in qNSCs is much less clear, with the same studies finding no changes to a mild increase in the total number of type-B cells (Gil-Perotin *et al*., 2006; Meletis *et al*., 2006). However, the likely compensatory effects of constitutive p53 deletion confound interpretation of these results and preclude assessment of adult-specific roles. Here, we combined a conditional and inducible p53 knock-out mouse model (*p53^icKO^*) with mechanistic *in vitro* assays to examine the function of p53 and its effectors in qNSCs of the adult SVZ. We found that p53 maintains NSC quiescence through regulation of FAO, independent of canonical p21 regulation. Strikingly, a fish-oil supplemented diet is sufficient to suppress the aberrant activation of qNSCs that results from p53 loss and in a somatic GBM model significantly delays glioblastoma development through restoration of FAO. Together, these findings show that dietary intervention can maintain the normal function of mutation-bearing tissues, highlighting potential strategies for cancer prevention.

## Results

### p53 enforces NSC quiescence

*p53^icKO^* mice were generated by crossing *p53^LoxP/LoxP^* or *p53^+/+^* mice to GLAST::CreERT2 animals and *p53* recombination was induced at 7 weeks of age using tamoxifen. A tdTomato (tdTom) reporter allele was also included in both backgrounds to trace recombined cells and their immediate progeny. Immunofluorescence analysis of primary NSCs acutely isolated from the SVZ 24h after tamoxifen administration confirmed efficiency of *p53* recombination in the tdTom^+^ population (>60%, Figures S1A and S1B). Furthermore, analysis of the neurogenic lineage by FACS, indicated that tdTom recombination was largely restricted to type-B cells, enabling assessment of acute effects of p53 loss on qNSCs, independent of known confounding effects on type-C and type-A progenitors (Figures S1C and S1D) (Gil-Perotin *et al*., 2006; Meletis *et al*., 2006).

We therefore investigated effects of p53 loss in both type-B cells that had previously activated and returned to quiescence (resting qNSCs) and type-B cells that had not yet activated (dormant qNSCs), which together constitute the two main subpopulations of SVZ NSCs (Obernier *et al*., 2018). For this, we carried out two complementary EdU-label retention experiments. To examine resting qNSCs, EdU was administered in the drinking water for 7 days, followed by a 5-day tamoxifen administration to recombine p53, and analysis of the entire SVZ in wholemount preparations 3 days later (Figure 1A). This protocol enables identification of resting qNSCs by labelling type-B cells that incorporated EdU during the 7-day pulse and then re-entered quiescence during the 8-day chase period, thereby appearing as EdU^+^/Ki67^-^/tdTom^+^ cells with radial morphology. We found a marked reduction in the percentage of resting qNSCs in *p53^icKO^* mice relative to controls, indicative of premature re-activation upon *p53* loss (Figure 1B and 1C). This was accompanied by an increase in pairs of rounded EdU^-^/Ki67^+^ and Ascl1^+^/td-Tom^+^ type-C cells, which likely represent the immediate progeny of non-label-retaining recombined type-B cells (Figures 1D, 1E; Figures S1E and S1F) (Obernier *et al*., 2018). In contrast, numbers of clusters of 3 and 4 cells detected in control and *p53^icKO^* animals were similar (Figures S1G and S1H), suggesting that observed phenotypes result from a direct effect of p53 loss in type-B cells rather than an increase in type-C proliferation. To examine effects on dormant qNSCs, we extended the EdU administration to 14 days, a time-window during which the vast majority of resting qNSCs incorporate EdU, whilst dormant qNSCs remain unlabelled (Obernier *et al*., 2018). p53 recombination was induced during the last 5 days of the EdU labelling period, and SVZ wholemounts analysed 1 day later (Figure 1F). This enables identification of dormant qNSCs that activate during the 1-day chase period as radial EdU^-^/Ki67^+^/tdTom^+^ cells. Our results revealed that p53 loss also results in premature activation of dormant qNSCs, as judged by an increase in EdU^-^/Ki67^+^/tdTom^+^ type-B cells (Figure 1G and 1H). This was accompanied by a trend towards an increase in EdU^-^/Ki67^+^ type-C cell pairs, which did not reach significance (Figure 1I). In addition, we detected a significant increase in EdU^+^/Ki67^+^ type C cell pairs, which likely represent immediate progenitors of recombined qNSCs (both resting and dormant) that activated during the tamoxifen/EdU administration period (Figure 1J). Thus, p53 maintains type-B cell quiescence and its deletion drives aberrant activation of both resting and dormant qNSCs.

**Figure 1.**
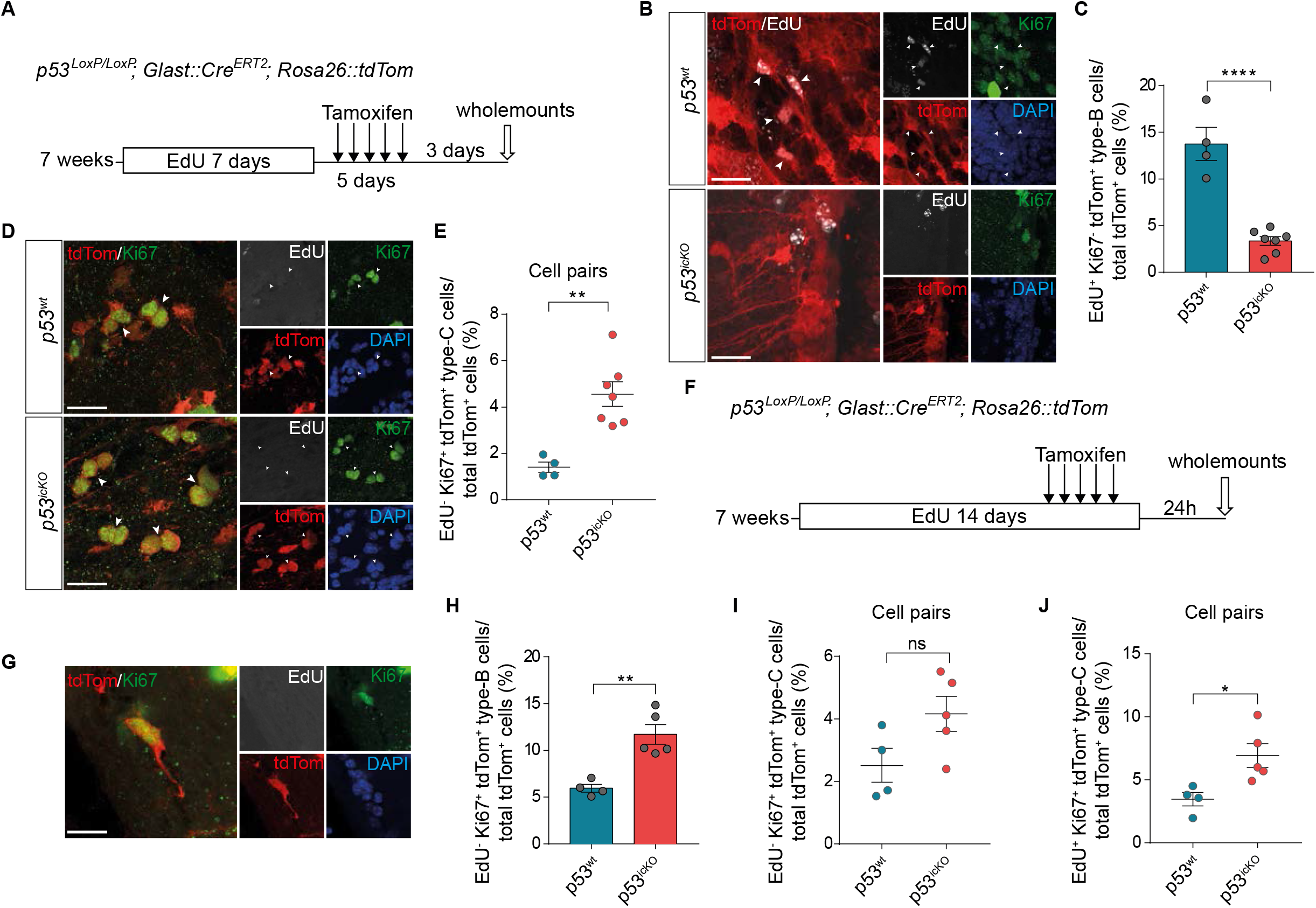
p53 controls NSC quiescence. **A,** Schematic of experimental outline. 7 weeks-old *p53^wt^* and *p53^icKO^* mice received EdU in the drinking water *ad libitum* for 7 days, followed by Tamoxifen injection for 5 consecutive days. V-SVZ wholemount explants were collected 3 days after the last injection and processed. **B,** Representative confocal images of EdU^+^ (grey)/Ki67^-^ (green)/tdTom^+^ (red) label-retaining resting type-B cells (white arrowheads) in *p53^wt^* and *p53^icKO^* SVZ wholemounts. Nuclei are counterstained with DAPI (blue). Scale bar=20μm. **C,** Quantification of resting type-B cells in both *p53^wt^* and *p53^icKO^* genotypes shown in B. Bar plots represent percentages of EdU^+^/Ki67^-^/tdTom^+^ cells over the total number of tdTom^+^ type-B and type-C cells. Mean±SEM, *p53^wt^* n=4, *p53^icKO^* n=7, ****p<0.0001, unpaired two-tailed Student t-test. **D,** Representative fluorescence images of EdU^-^ (grey)/Ki67^+^ (green)/tdTom^+^ (red) pairs of type-C cells (white arrowheads) in *p53^wt^* and *p53^icKO^* wholemounts. Nuclei are counterstained with DAPI (blue). Actively proliferating cells are positive for Ki67 staining (green). Scale bar=20μm. **E,** Quantification of EdU^-^/Ki67^+^/tdTom^+^ type-C cell pairs in both genotypes. Data are shown as percentages over the total number of tdTom^+^ type-B and type-C cells. Mean±SEM, *p53^wt^* n=4, *p53^icKO^* n=7, **p<0.01, unpaired twotailed Student t-test. **F,** Schematic of experimental outline. 7 weeks-old *p53^wt^* and *p53^icKO^* mice received EdU in drinking water *ad libitum* for 14 days and were injected with Tamoxifen over the last 5 days of EdU administration. V-SVZ wholemount explants were collected 24h after the last injection and stained. **G,** Example of a dormant type-B cell that has undergone activation as judged by presence of Ki67 and lack of EdU labelling. Scale bar=20μm. **H,** Quantification of percentage of EdU^-^/Ki67^+^/tdTom^+^ active type-B cells in both *p53^wt^* and *p53^icKO^* genotypes over the total number of type-B and type-C tdTom^+^ cells. Mean±SEM, *p53^wt^* n=4, *p53^icKO^* n=5, **p<0.01, unpaired two-tailed Student t-test. **I-J,** Quantification of EdU^-^ /Ki67^+^/tdTom^+^ (I) and EdU^+^/Ki67^+^/tdTom^+^ (J) pairs of type-C cells in *p53^wt^* and *p53^icKO^* brains.Percentage over the total number of tdTom^+^ type-B and type-C cells is shown. Mean±SEM, *p53^wt^* n=4, *p53^icKO^* n=5, ns= not significant, *p<0.05, unpaired two-tailed Student t-test. See also Figure S1.

### p53 regulates Fatty Acid Oxidation in qNSCs

We next sought to identify the mechanisms by which p53 enforces quiescence. For this, we exploited a previously established co-culture assay in which neural stem/progenitor cells (NPCs) acquire phenotypic and molecular hallmarks of quiescence upon direct cell-cell contact with primary brain microvascular endothelial cells for 48h (bmvEC) (Ottone *et al*., 2014). To further assess the robustness of this model system, we profiled the transcriptomes of NPCs cultured alone or with bmvEC by RNA-sequencing (Supplemental Table S1). Comparison with published transcriptional signatures of qNSCs acutely purified from the SVZ niche (Codega *et al*., 2014; Llorens-Bobadilla *et al*., 2015; Mizrak *et al*., 2019), revealed that endothelial-induced quiescence mimics *in vivo* phenotypes to a significant extent, with an overlap of 384 upregulated (25.1% overlap with *in vivo* qNSC signature, p=3.7*10^−109^) and 246 downregulated genes (36.4% overlap with *in vivo* aNSC signature, p=1.1*10^−131^, Figure S2A). In addition, KEGG and GO slim analysis showed that several quiescence-specific biological processes are reproduced in co-culture, including upregulation of type-B fate markers (transporter activity, cell differentiation, glutamatergic and cholinergic synapse), increased adhesion and signalling (cell adhesion molecules, signalling activity), increased lipid metabolism (lipid metabolic process) and decreased cell proliferation (cell-cycle, DNA replication, cell population proliferation) (Figures S2B and S2C) (Codega *et al*., 2014; Tong *et al*., 2014; Llorens-Bobadilla *et al*., 2015; Paul, Chaker and Doetsch, 2017; Mizrak *et al*., 2019). We next performed functional studies to assess whether endothelial-induced quiescence *in vitro* is p53-dependent as it is *in vivo*. Luciferase reporter assays revealed that p53 activity was increased in NPCs cultured with bmvECs compared to NPCs cultured alone (Figure 2A). Furthermore, primary *p53^LoxP/LoxP^* NPCs acutely recombined with Adeno-Cre viruses (*p53^-/-^* NPCs) underwent a less pronounced cell-cycle arrest on bmvEC than Adeno-Control-transduced controls (*p53^+/+^* NPCs), consistent with previous reports (Figure 2B; Figures S2D and S2E) (Mathieu *et al*., 2008). Together, these data suggest that the co-culture system reflects *in vivo* quiescence phenotypes and can be used to identify p53 effectors.

**Figure 2.**
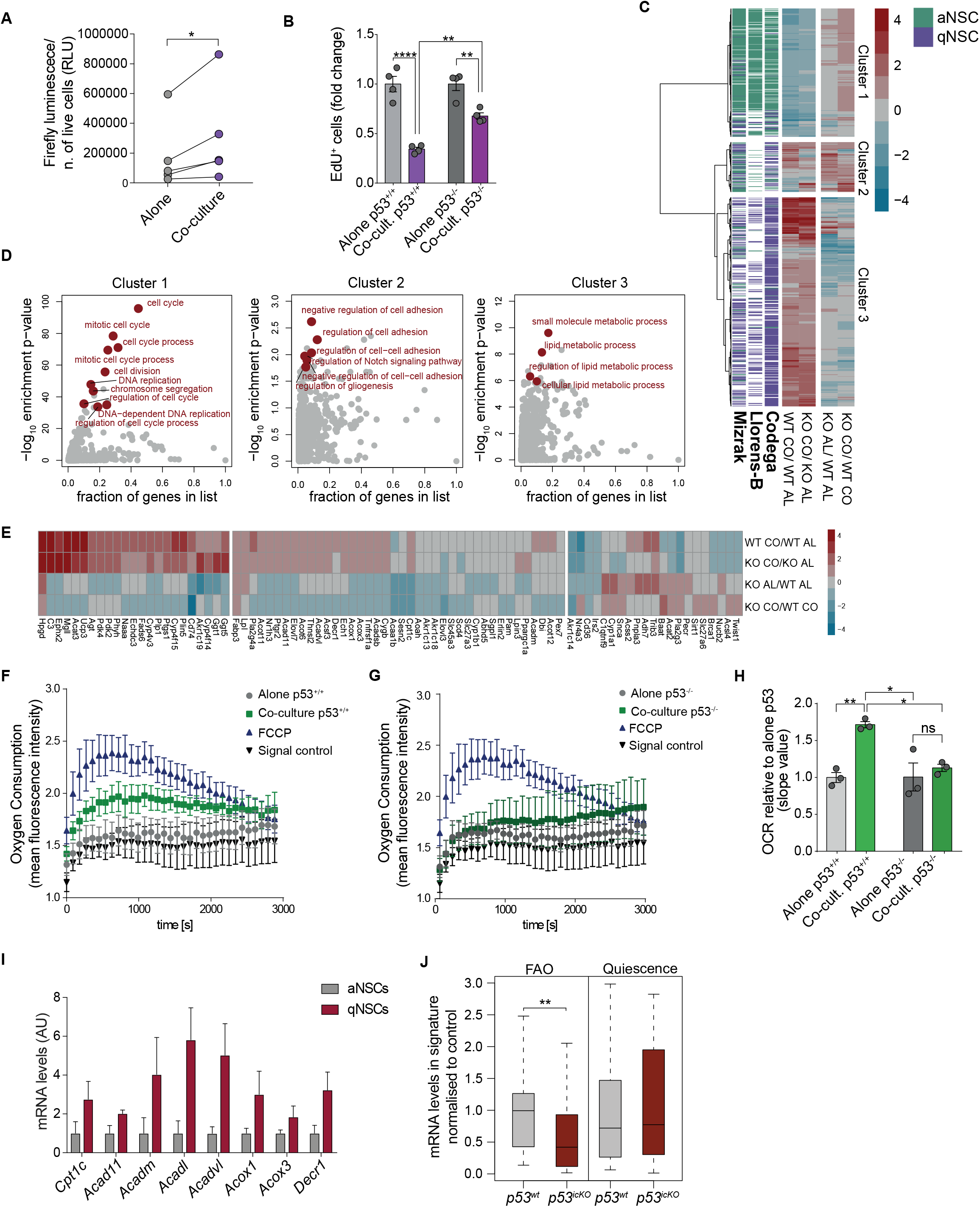
p53 regulates FAO in qNSCs. **A,** p53 promoter activity of NPCs cultured alone or with primary brain microvascular endothelial cells (bmvEC; Co-culture) for 48h. Luciferase intensity normalized to viable cell number is shown. Data are represented as Relative Light Units (RLU). n=5 independent cultures. *p<0.05, paired two-tailed Student t-test. **B,** Quantification of EdU FACS profiles of *p53^+/+^* and *p53^-/-^* NPCs cultured alone or with bmvEC for 48h and pulsed for 2h with EdU. For each genotype, fold change relative to its own alone control is shown. Mean±SEM, n=4 independent experiments, **p<0.01, ****p<0.0001, Two-way ANOVA with Tukey’s multiple comparisons test. **C,** Hierarchical clustering of RNA-seq log_2_ expression ratios between of *p53^+/+^* (WT) and *p53^-/-^* (KO) NPCs in alone (AL) and co-culture (CO) conditions alongside published data-sets from qNSCs and aNSCs acutely FACS purified from the SVZ of adult mouse brains (Codega *et al*., 2014; Llorens-Bobadilla *et al*., 2015; Mizrak *et al*., 2019). Cluster 1: genes upregulated in p53^-/-^ NPCs/bmvEC co-cultures, Cluster 2: set of genes that do not change in both in alone and in co-culture conditions. Cluster 3: genes downregulated in either p53^-/-^ NPCs alone or in co-culture. **D,** GO term enrichment analysis of the three clusters from C. Enrichment p-values are shown as a function of the fraction of genes from each cluster present in highlighted GO categories. **E,** Hierarchical clustering of RNA-seq log_2_ expression ratios for genes involved in lipid metabolism in *p53^+/+^* (WT) and *p53^-/-^* (KO) NPCs in alone (AL) and in co-culture (CO) conditions. **F-G,** FA-driven oxygen consumption over time of *p53^+/+^* (F) and *p53^-/-^* (G) NPCs alone and in co-culture with bmvECs plotted as mean fluorescent intensity of n=3 independent experiments. FCCP treatment was used as positive control and background fluorescence signal as negative control (absence of cell respiration). **H,** Quantification of the oxygen consumption rate (OCR) of *p53^+/+^* and *p53^-/-^* NPCs cultured alone or with primary endothelial cells. OCR is represented as the slope value of the linear proportion of the signal profile shown in F and G. Mean±SEM, n=3 independent experiments, ns=not significant, *p<0.05, **p<0.01, Two-way ANOVA with Tukey’s multiple comparisons test. **I,** Quantitative RT–PCR analysis of indicated FAO genes in aNSCs and qNSCs acutely FACS-purified from the SVZ of GFAP::GFP mice. Mean±SEM, n=3 independent sorts. **J,** qRT–PCR analysis of FAO and quiescence gene signatures in qNSCs acutely FACS-purified from the SVZ of *p53^wt^* (grey) and *p53^icKO^* (red) mice 24h following tamoxifen administration. Boxplots represent median, interquartile range, and most extreme data points that are not more than 1.5 times the interquartile range. n=4 independent sorts. p=0.0085 for FAO signature, p=0.98 for quiescence signature, two-sided Wilcoxon test. See also Figure S2.

To do so, we characterised the transcriptional programmes controlled by p53 using the same adenoviral-mediated acute loss of function approach as above. *p53^+/+^* and *p53^-/-^* NPCs were cultured alone or on endothelial monolayers for 48h and subjected to RNA-sequencing (Supplemental Table S1). Hierarchical clustering of the genes shared between NPCs in coculture and qNSCs *in vivo* (Figure S2A), identified three groups of genes based on expression changes in *p53^-/-^* NPCs (Figures 2C and 2D; Supplemental Table S1). These corresponded to genes that did not significantly change (cluster 2), increased (cluster 1) or decreased (cluster 3) in expression upon p53 loss. Cluster 2 comprised genes related to type-B cell identity, including glial markers, adhesion and Notch signalling, suggesting that p53 does not directly control type-B fate (Codega *et al*., 2014; Llorens-Bobadilla *et al*., 2015). In agreement with this, immunostaining of *p53^+/+^* and *p53^-/-^* NPCs co-cultures confirmed that GFAP expression and neurosphere-like morphology were unaltered upon p53 loss (Figure S2E). Cluster 1 predominantly included genes involved in cell-cycle, as expected from the increased proliferation of *p53^-/-^* cultures (Figure 2B). Cluster 3 was highly enriched in signatures of lipid metabolism, suggesting that p53 may control the metabolic profile of qNSCs.

In response to stress or DNA damage, p53 mediates cell-cycle arrest predominantly through transcriptional activation of *p21/Cdkn1a* (Bieging, Mello and Attardi, 2014; Engeland, 2018). We therefore assessed p21 function in our system, by comparing the response of wildtype (*p21^+/+^*) and p21 knock-out (*p21^-/-^*) NPCs to endothelial co-culture. Surprisingly, we found that *p21^-/-^* cells arrested to the same extent as *p21^+/+^* cells (Figure S2F). Consistent with this, p53 binding to the *p21* promoter and *p21* mRNA levels were similar in wild type NPCs cultured alone and with endothelial cells (Figure 3E; Figure S2G). Thus, p53-induced NPC quiescence is p21-independent.

**Figure 3.**
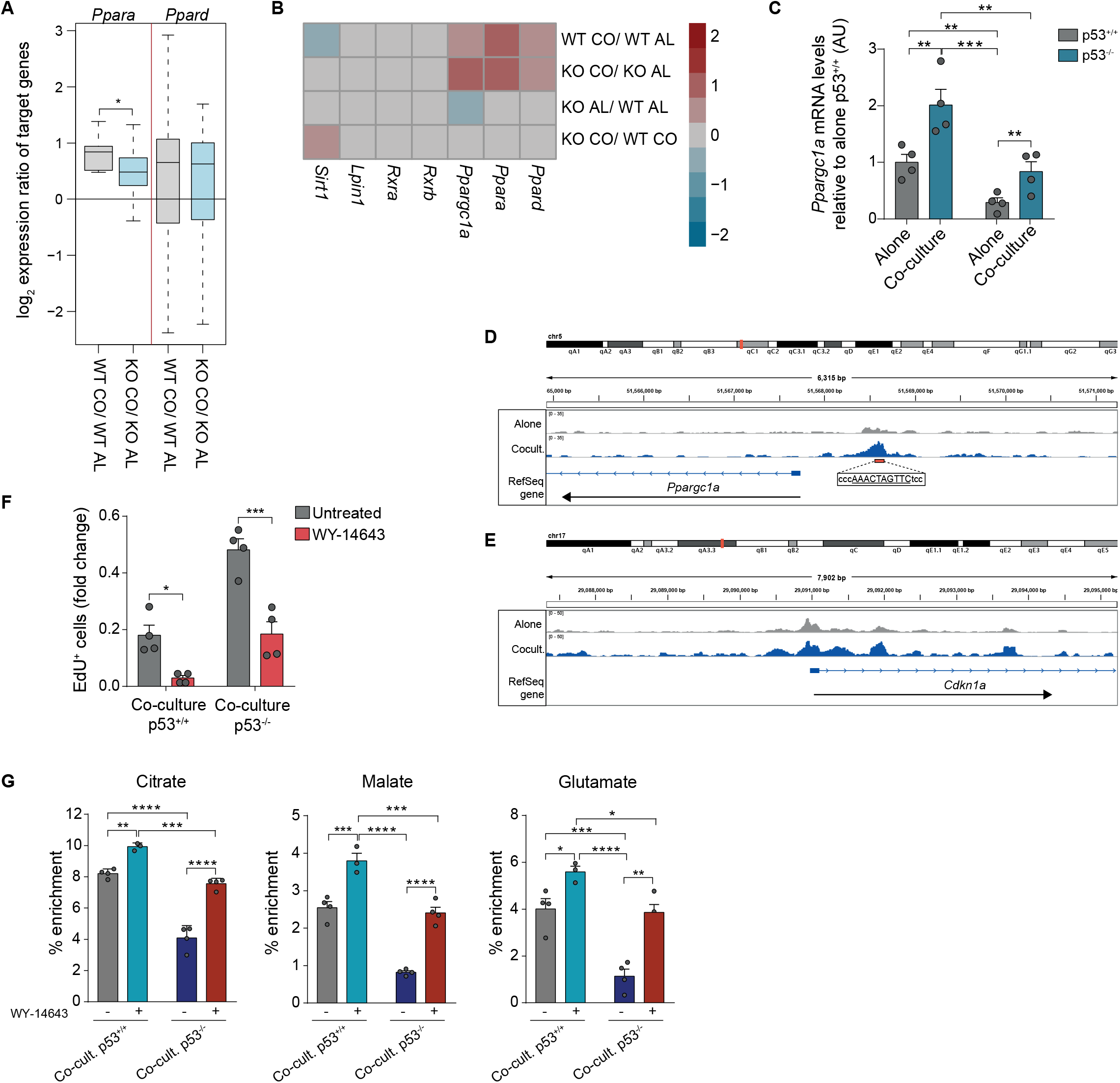
p53 mediates quiescence through PPARα. **A,** RNA-Seq log_2_ expression ratios of *Ppara* and *Ppard* target genes in *p53^+/+^* (WT) and *p53^-/-^* (KO) NPCs alone (AL) and in coculture (CO), p=0.037, two-sided Wilcoxon test. **B,** RNA-seq log_2_ expression ratios of FAO regulators in *p53^+/+^* (WT) and *p53^-/-^* (KO) NPCs alone (AL) and in co-culture (CO). **C,** Quantitative RT–PCR analysis of *Ppargc1a* mRNA levels in *p53^+/+^* and *p53^-/-^* NPCs alone and in co-culture. Fold changes relative to alone *p53^+/+^* are shown. Mean±SEM, n=4 independent experiments, **p<0.01, ***p<0.001, Two-way ANOVA with Tukey’s multiple comparisons test. **D-E,** Read coverage for p53 CUT&RUN experiments at *Ppargc1a* (D) and *Cdkn1a* (E) gene loci for NPCs alone (grey tracks) and in co-culture (blue tracks). The red box depicts the p53 responsive element identified upstream the TSS of *Ppargc1a*. **F,** FACS quantification of EdU^+^ *p53^+/+^* and *p53^-/-^* NPCs in co-culture, before and after treatment with the PPARα agonist WY-14643 for 24h. Fold change to respective alone controls is shown. Mean±SEM, n=4 independent experiments, *p<0.05, ***p<0.001, Two-way ANOVA with Tukey’s multiple comparisons test. **G**, ^13^C enrichment following a 24h incubation with [U- ^13^C]-palmitic acid in *p53^+/+^* and *p53^-/-^* NPCs, in the presence or absence PPARα agonist WY-14643. Bar plots show the ^13^C-incorporation in citrate and malate (TCA intermediates) and also glutamate. Data are plotted as the % ^13^C enrichment for all isotopologues of each metabolite. Mean±SEM, n=4 replicates, *p<0.05, **p<0.01, ***p<0.001, ****p<0.0001, Two-way ANOVA with Sidak’s multiple comparisons test. See also Figure S3.

Fatty acid oxidation (FAO) has emerged as an important mediator of stem cell quiescence (Ito *et al*., 2012; Knobloch *et al*., 2017; Mihaylova *et al*., 2018; Wang *et al*., 2018). In the hippocampus, FAO was shown to maintain NSC quiescence and modulation of FAO levels resulted in changes in the balance between quiescent and proliferative states of NSCs (Knobloch *et al*., 2017; Wani *et al*., 2022). As we observed changes in lipid metabolism signatures in *p53^-/-^* NPCs/bmvEC co-cultures (Figure 2D), we next examined the role of FAO downstream of p53. We assessed expression of FAO genes in the RNA-seq data and found that many enzymes involved in FA metabolism were deregulated in *p53^-/-^* relative to *p53^+/+^* NPCs (Figure 2E). To determine whether these expression changes were also accompanied by functional changes, we first measured FA-driven oxygen consumption in *p53^+/+^* and *p53^-/-^* NPCs cultured alone and with bmvECs. While *p53^+/+^* NPCs increased FAO upon endothelial co-culture (Figures 2F and 2H), *p53^-/-^* NPCs did not (Figures 2G and 2H). Interestingly, despite FAO signatures being downregulated in both proliferating and quiescent *p53^-/-^* NPCs, only in co-culture did this result into a change in metabolic state, confirming that a shift to FAO is a quiescence-specific phenotype (Knobloch *et al*., 2017).

Next, we asked whether p53 regulates the FAO transcriptional programme *in vivo*. First, we acutely FACS-purified quiescent and active type-B cells from the SVZ of GFAP::GFP mice (identified as GFP^+^/CD24^-^/CD133^+^/EGFR^-^ and GFP^+^/CD24^-^/CD133^+^/EGFR^+^, respectively) and used qRT-PCR to measure expression of a panel of FAO genes found to be regulated by p53 in Figure 2E (see also Supplemental Table S1). We found that expression levels were higher in qNSCs relative to aNSCs, consistent with increased lipid metabolism in quiescent cells (Figure 2I; Figures S2H and S2I) (Codega *et al*., 2014; Llorens-Bobadilla *et al*., 2015). Furthermore, analysis of qNSCs acutely FACS-purified from *p53^icKO^* mice 24h post-recombination (identified as tdTom^+^/CD24^-^/CD133^+^/EGFR^-^, Figure S1C) revealed an overall downregulation of the FAO gene signature upon *p53* deletion, in the absence of changes in quiescence markers, confirming that p53 selectively controls the FAO programme *in vitro* and *in vivo* (Figure 2J).

### p53 mediates quiescence through PPARα

Our results thus far suggest that p53 may maintain qNSCs by enforcing the lipid metabolic programme that sustains quiescence through transcriptional regulation of FAO pathway genes. In qNSCs of the hippocampal neurogenic niche, FAO is partially controlled by peroxisome proliferator-activated receptor alpha (PPARα), a ligand-activated transcription factor and master regulator of FAO genes (Knobloch *et al*., 2017). As the vast majority of FAO genes found to be dysregulated in *p53^-/-^* NPCs (Figure 2E) are known PPAR targets (Fang *et al*., 2016), we hypothesised that PPARs may also mediate p53 effects. Both *Ppara* and *Ppard* isoforms were found to be expressed in NPCs and upregulated in co-culture (Figure 3B). However, while expression of target genes of both isoforms increased in quiescent NPCs, only *Ppara* targets were p53-dependent, suggestive of a p53/PPARα crosstalk (Figure 3A). To identify the underlying mechanisms, we examined expression of PPARs themselves, alongside their key regulators (Sugden, Caton and Holness, 2010), in the RNA-seq data. While most genes were not affected by p53 loss, expression of the PPARα co-activator Peroxisome proliferator-activated receptor gamma coactivator 1-alpha (*Ppargc1a/Pgc1a*) (Sugden, Caton and Holness, 2010), was strongly downregulated in *p53^-/-^* NPCs cultured alone (Figures 3B and 3C). In the RNA-seq data we only observed a modest reduction in *Ppargc1* levels in *p53^-/-^* cocultures relative to *p53^+/+^* co-cultures, which did not reach significance (Figure 3B). This was likely due to the relatively small overall fold change in *Ppargc1* transcript upon quiescence and the variability of the biological replicates. However, subsequent qPCR validation experiments indicated that *Ppargc1* levels decreased significantly also in *p53^-/-^* NPCs cocultures, reverting to the basal levels of *p53^+/+^* NPCs cultured alone (Figure 3C). Interestingly, despite *Ppargc1* expression being overall lower in *p53^-/-^* than in *p53^+/+^* NPCs, *Ppargc1* levels still increased in *p53^-/-^* co-cultures, suggestive of additional p53-independent mechanisms (Fernandez-Marcos and Auwerx, 2011). Furthermore, CUT&RUN analysis of wildtype NPCs before and after co-culture with endothelial cells revealed direct binding of p53 at a response element within the *Ppargc1a* promoter, specifically upon co-culture-induced quiescence (Figure 3D). Indeed, *Ppargc1a* was one of the genes identified to be both bound by p53 in the CUT&RUN data and deregulated upon p53 loss in the RNA-sequencing data (Figure S3A). Together, these results point to a mechanism whereby elevated p53 activity in qNSCs induces *Ppargc1a* transcription. In turn, PPARGC1a enhances PPARα activity to induce transcription of lipid catabolic enzymes resulting in increased FAO and quiescence (Figure 4L). To functionally test this model, we asked whether increasing PPARα activity through administration of exogenous ligands would compensate for the decrease of PPARGC1a in *p53^-/-^* NSCs and rescue quiescence (Vega, Huss and Kelly, 2000). Treatment with the PPARα agonist WY-14643 increased expression of FAO genes and restored the cell-cycle arrest of *p53^-/-^* NPCs in co-culture to the same levels as *p53^+/+^* NPCs, without affecting stemness (Figure 3F; Figures S3B, S3C and S3D). The rescue was dependent on PPARα and not caused by off target effects of WY-14643 because it was lost in *Ppara* knock-out cells (Figures S3E; Figures S4F and S4G). To further confirm that the effects of WY-14643 were mediated by FAO downstream of PPARα, we exposed quiescent *p53^+/+^* and *p53^-/-^* NPCs to ^13^C-palmitate in the presence or absence of WY-14643 and traced the incorporation of radiolabelled carbons into TCA intermediates and amino acids using mass spectrometry (GC-MS). We found that while incorporation of ^13^C into both TCA intermediates and amino acids derived from TCA intermediates was significantly decreased in untreated *p53^-/-^* co-cultures, as expected, WY-14643 treatment restored it to the levels of *p53^+/+^* cocultures (Figure 3G; Figure S3F). Together, these experiments indicate that p53 maintains NSC quiescence through regulation of FAO via a PPARGC1a/ PPARα axis.

**Figure 4.**
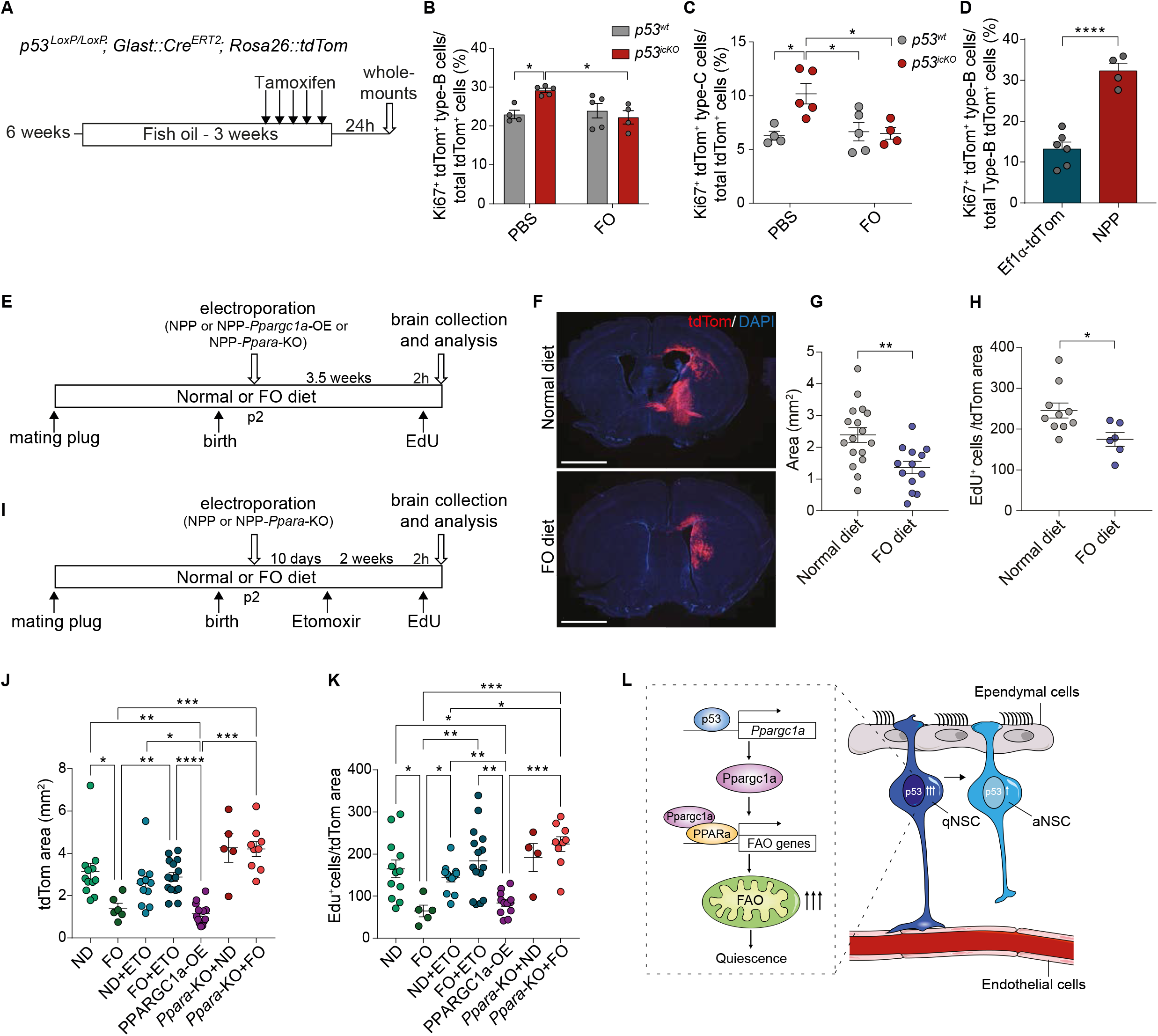
A fish oil supplemented diet delays tumour initiation. **A,** Schematic of experimental outline. 6 weeks-old *p53^wt^* and *p53^icKO^* mice received omega-3 enriched fish oil via oral gavage for 3 weeks and were injected with Tamoxifen over the last 5 days of fish oil treatment. V-SVZ wholemount explants were collected 1 day after the last injection and stained for analysis. **B-C,** Quantification of percentage of Ki67^+^/tdTom^+^ active type-B cells (B) and of Ki67^+^/tdTom^+^ type-C cell pairs (C) in the SVZ of *p53^wt^* and *p53^icKO^* after administration of fish oil or PBS as control. Data are represented as percentage over the total number of tdTom^+^ type-B and type-C cells. Mean±SEM, *p53^wt^* + PBS n=4, *p53^icKO^* + PBS n=5, *p53^wt^* + Fish oil n=5, *p53^icKO^* + Fish oil n=4. *p<0.05, Two-way ANOVA with Tukey’s multiple comparison test. **D**, Quantification of Ki67^+^/tdTom^+^ active type-B cells in SVZ wholemounts of C57BL/6 mice 10 days after electroporation of EF1α-tdTomato or NPP (*Nf1,Pten,Trp53*) plasmids into the SVZ. Mean±SEM, EF1α-tdTomato n=6, NPP n=4. ****p<0.0001, unpaired two-tailed Student t-test. **E**, Schematic of experimental outline. Pregnant females were placed on fish oil supplemented diet or normal diet from conception and pups electroporated at P2. Early tumours were collected after 3.5 weeks following a 2-hour EdU pulse. **F-G**, Representative fluorescence images (F) and quantifications (G) of tdTomato-labelled tumours in mice fed normal or fish oil (FO)-supplemented chow. Scale bar=2mm. Mean±SEM, normal diet n=17, FO n=13, **p<0.01, unpaired two-tailed Student t-test. **H**, Quantification of the number of EdU^+^ cells/tdTom^+^ tumour area in the bulks of gliomas developing in the brains of mice fed control and FO-supplemented diet. Data are represented as the number of EdU^+^ nuclei normalized to the dtTom^+^ tumour area. Mean±SEM, normal diet n=10, FO n=6, *p<0.05, unpaired two-tailed Student t-test. **I**, Schematic of experimental outline. Treatment was as in E with the exception that 10 days after electroporation at P2, pups received etomoxir (ETO) for 2 weeks. **J-K**, Quantifications of tdTomato-labelled tumour size (J) and of the number of EdU^+^ cells/tdTom^+^ tumour cells (K) in brains of mice fed a normal (ND) or fish oil-supplemented (FO) diet upon treatment with etomoxir (ETO). tdTomato^+^ tumour area and number of EdU^+^ cells/tdTom^+^ tumour cells were also assessed in NPP-*Ppargc1a*-OE (PPARGC1a-OE) fed a ND and in NPP-*Ppara*-KO tumours fed ND or FO diets (*Ppara* KO+ND and *Ppara* KO+FO). Mean±SEM, ND n=12, FO n=6, ND+ETO n=11, FO+ETO n=14, PPARGC1a-OE n=13, *Ppara* KO+ND n=5, *Ppara* KO+FO n=9, *p<0.05, **p<0.01, ***p<0.001, ****p<0.0001, Ordinary One-way ANOVA with Dunnett’s T3 multiple comparisons test. **L,** Model of p53 regulation of NSCs quiescence in the SVZ. See also Figure S4.

### Dietary fish oil supplementation delays tumour initiation

Our results suggest that loss of NSC quiescence via disruption of their metabolic state may be a key mechanism by which p53 mutations drive GBM. We therefore hypothesised that restoring PPARα-dependent FAO downstream of mutant p53 may reverse these effects and ultimately suppress tumourigenesis. PPARα is a nutrient sensor and can be activated by dietary polyunsaturated FAs (PUFAs), providing a potential strategy for tumour prevention through diet (Contreras, Torres and Tovar, 2013). To test this strategy directly, we examined effects of dietary supplementation with fish oil, a natural source of Omega-3 PUFAs, on p53 phenotypes *in vivo*. Fish oil, or PBS control, was administered by gavage to *p53^wt^* or *p53^icKO^* mice for a total of 3-weeks during the last 5 days of which the animals also received tamoxifen prior to analysis of the SVZ 24h later (Figure 4A). Strikingly, dietary supplementation with fish oil fully reversed the effect of acute p53 loss in qNSCs *p53^icKO^*, suppressing the increase in both activated type-B and early type-C progenitors compared to vehicle-treated animals (Figures 4B and 4C). Thus, diet alone is sufficient to maintain quiescence of p53-deficient NSCs.

To determine whether diet could also impact oncogenic transformation, we examined effects of fish oil supplementation on tumour initiation from NSCs in a somatic mouse model of GBM based on CRISPR/Cas9 gene editing and PiggyBac transposition technology (Zuckermann *et al*., 2015; Brooks *et al*., 2021; Garcia-Diaz *et al*., 2021). The model is driven by combined inactivation of the tumour suppressor genes *Nf1, Pten and Trp53* (hereon NPP model) in endogenous postnatal SVZ NSCs and includes stable integration of a tdTomato reporter gene in all tumour cells (Figure S4A). A non-integrating plasmid encoding for the PiggyBase transposase and Cas9 under the control of a NSC-specific promoter alongside an integrating PiggyBac vector, carrying the CRISPR guides to tumour suppressors and tdTomato, are coelectroporated into the lateral ventricles of wild-type P2 pups. Upon electroporation, transient Cas9/sgRNA expression results in inactivation of *Nf1, Pten and Trp53*, whereas PiggyBase-mediated integration of the PiggyBac vector ensures stable expression of the oncogenes and the td-Tomato reporter in the targeted NSCs and their progeny (Figure S4A). As in genetically engineered mouse models carrying the same mutations (Kwon *et al*., 2008; Alcantara Llaguno *et al*., 2009), the NPP model gives rise to brain tumours with histological and molecular features of GBM at high penetrance (Brooks *et al*., 2021; Garcia-Diaz *et al*., 2021). Importantly, the introduction of the three mutations resulted in a robust increase in the percentage of NSCs undergoing activation at 10 days post-electroporation, relative to control mice electroporated with a control plasmid expressing tdTomato alone (Figure 4D). This indicates that aberrant activation of qNSCs is an early event in tumour initiation and that the effects of acute p53 loss on qNSCs are similar whether it occurs alone or in the tumorigenic context of *Nf1* and *Pten* inactivation. Females were fed a high fish oil or control diet from conception and throughout pregnancy and lactation. Tumourigenesis was then induced in the pups at postnatal day 2 and brain tissue analysed 3.5 weeks later, a time at which early lesions can be detected in control animals (Figures 4E and 4F). Remarkably, analysis of resulting lesions revealed that they were significantly smaller and less proliferative in the high fish oil diet-fed group relative to controls, as judged by measurement of td-Tomato^+^ area and EdU incorporation, respectively (Figures 4G and 4H). We next carried out gain- and loss-of-function experiments to determine whether dietary fish oil administration suppresses tumour initiation *in vivo* through the same p53-regulated mechanism we identified *in vitro*. To determine the contribution of FAO downstream of fish oil administration, we repeated the protocol described above in the presence or absence of etomoxir (Figure 4I) and found that FAO inhibition completely abrogated fish oil effects, leading to the development of larger and more proliferative lesions, similar to early NPP control tumours (Figures 4J and 4K). Furthermore, quantification of the proportion of quiescent and active NSCs in the four tumour groups, revealed that fish oil suppressed the aberrant NSC activation seen in control or etomoxir-treated mice fed a control diet, reverting it to normal neurogenesis levels (see Ef1α-tdTom in Figure 4D). In contrast, etomoxir abrogated fish oil effects, restoring NSC activation to normal diet levels (Figure S4B). These experiments confirm that fish oil maintains mutation-bearing NSCs in a quiescent state and suppresses tumour initiation through FAO.

Furthermore, we assessed whether FAO induction downstream of fish oil depends on the identified PPARGC1a/PPARα axis. The NPP construct was therefore modified to incorporate the *Ppargc1a* gene (Figure S4A), resulting in its overexpression and induction of p53-regulated FAO genes in the tumour cells (Figures S4C, S4D and S4E). Ensuing early tumours in animals fed a normal diet were significantly smaller and less proliferative than controls, closely phenocopying fish oil effects (Figures 4J and 4K). We next deleted the *Ppara* gene by introduction of a CRISPR sgRNA to *Ppara* into the NPP construct (Figure S4A) and, upon confirmation of successful knock-out of the gene (Figures S4F and S4G), used this construct to initiate tumours in animals fed a normal or high fish oil diet. Remarkably, the *Ppara* KO tumours reverted to control size despite dietary fish oil administration (Figures 4J and 4K), confirming that fish oil suppresses initiation through PPARα. We conclude that diet counteracts tumour-initiating mutations to delay gliomagenesis.

## Discussion

Mounting evidence suggests that normal tissues commonly bear a variety of genetic changes, including driver mutations (Bissell and Hines, 2011; Martincorena *et al*., 2017, 2018). How these mutations remain silent despite their tumour-initiating potential, is a fundamental and still unresolved question in cancer biology. It has been proposed that a normal tissue microenvironment acts as a tumour suppressive mechanism by dominantly keeping mutations in check, for example via dampening of mitogenic signalling pathways or ECM-tumour cell interactions (Pasquale, 2010; Bissell and Hines, 2011). Our study identifies diet as a key contributing factor in mutation silencing.

Mutations in p53 are among a handful of genetic changes that have been found to be shared between tumour-free SVZ and matching tumours of GBM patients (Lee *et al*., 2018). This important discovery points to NSCs as GBM cells of origin and to p53 as a potential tumour-initiating mutation. Although only approximately 30% of mutations in the p53 gene are inactivating, with the rest being gain-of-function mutations, p53 is also inactivated in tumours with MDM1/2/4 amplifications (~15% of GBMs) and deletion of the *Cdkn2a* locus (~60% of GBMs). Thus, p53 loss-of-function is a common and important event in the aetiology of the disease (Brennan *et al*., 2013; Lee *et al*., 2018). Consistent with this, p53 has been previously linked to the regulation of adult neurogenesis. However, as previous studies have been restricted to constitutive p53 knock-out mouse models, they could not inform on the specific impact of p53 inactivation on NSC biology (Gil-Perotin *et al*., 2006; Meletis *et al*., 2006; Lee *et al*., 2018). By inducing temporally-controlled recombination of p53 selectively in quiescent NSCs, we found that p53 loss skews the homeostatic balance between quiescence and activation of type-B cells. In the context of additional mutations, aberrant NSC activation was an early event in tumour initiation. This is consistent with observations in other niches in which driver mutations have been shown to initiate tumourigenesis by prematurely activating quiescent stem cells (Zhang *et al*., 2006; He *et al*., 2007; Westphalen *et al*., 2014; White *et al*., 2014; Moon *et al*., 2017).

We found that p53 regulation of quiescence is independent of its canonical effector p21. This is unexpected given the major role of p21 in blocking proliferation via the DREAM complex (Engeland, 2018) and underscores the fundamental differences in p53-dependent regulatory mechanisms between homeostasis and response to stress (Maddocks and Vousden, 2011). Instead, we show that p53 maintains NSC quiescence through regulation of FAO, highlighting the dominant role of the cellular metabolic state in directing decisions between quiescence and activation, consistent with previous studies (Ito *et al*., 2012; Knobloch *et al*., 2017; Mihaylova *et al*., 2018; Wang *et al*., 2018).

Interestingly, unlike in other systems where p53 directly controls transcription of FAO genes (Bensaad *et al*., 2006; Jiang *et al*., 2015), our data reveal that in quiescent NSCs p53 induces FAO via a PPARGC1a/PPARα axis. There is a precedent for a bidirectional crosstalk between p53 and PPARGC1a in cancer, where p53 was shown to induce PPARGC1a expression and PPARGC1a to control p53-dependent cell fate decisions in response to oxidative or metabolic stress, respectively (Sen, Satija and Das, 2011; Aquilano *et al*., 2013). Similarly, PPARGC1a was shown to activate FAO via PPARα downstream of PML in breast cancer (Carracedo *et al*., 2012). It is of note that repression of PPARGC1a downstream of p53 has also been reported in hTERT deleted cells (Sahin *et al*., 2011). This apparent discrepancy is likely due to differences in the positioning of the p53 responsive element utilised, which was proximal (~900bp) to the TSS in our system, consistent with transactivation, and more distal (~2.8Kb) in the context of telomerase disfunction, consistent with repression (Sahin *et al*., 2011; Fischer, 2017). It is tempting to speculate that the p53/PPARGC1a/PPARα may represent a general homeostatic pathway in normal cells, which is highjacked in cancer to promote tumour growth by rewiring the cell metabolic state.

These findings have important therapeutic implications as we showed that, in the context of the NPP model, early GBM development was dramatically suppressed by fish oil. This suggests that dietary intervention may be an effective therapeutic strategy for suppressing tumour initiation. As quiescent stem cells often share a common metabolic profile and can act as cancer cell-of-origin across many tissues, dietary intervention may provide a more general approach for cancer prevention (Knobloch and Widmann, 2018).

## Acknowledgements

This work was funded by Cancer Research UK (S.P., H.S.R.), NIHR Biomedical Research Centre (V.A., M.C.) and the Medical Research Council (S.P., S.M., V.A., T.D., A.M.). It used the computing resources of the UK Medical Bioinformatics partnership (UK MED-BIO; aggregation, integration, visualization, and analysis of large, complex data), which is supported by the Medical Research Council and Imperial College High Performance Computing Service. We thank M. Goetz for GLAST::CreERT2 mice and O. Samson for *Cdkn1a-/-* tissue, J. Manji for help with microscopy, Y. Guo, G. Morrow and B. Wilbourn for help with FACS, S. Khadayate for help with bioinformatics, D. Helmlinger and L. Game for technical advice and F. Guillemot for helpful discussion and critical reading of the manuscript.

## Author Contributions

Conceptualization, S.P.; Methodology, V.A., T.D., C.G-D., A.G., H.K. and S.P.; Investigation, V.A., T.D., M.C., H.S.R., A.B, M.S.D.S., J.A.M, J.M. and S.P.; Formal analysis, A.S.M, H.S.R., M.S.D.S., J.M. and S.M.; Resources, S.M., A.P.G.; Writing – Original draft, V.A. and S.P.; Writing - Review and Editing V.A., S.P, S.M and A.P.G.; Visualization V.A., S.M., A.B and S.P.; Supervision, H.K., A.P.G, S.M. and S.P.; Funding acquisition, A.G, S.M., S.P..

## Methods

### Mice

All procedures were performed in compliance with the Animal Scientific Procedures Act, 1986 and approved by with the UCL Animal Welfare and Ethical Review Body (AWERB) in accordance with the International guidelines of the Home Office (UK). GFAP::GFP mice were obtained from The Jackson Laboratory (Jax 003257). *p53^icKO^* were generated by crossing GLAST::CreERT2 mice to animals carrying a loxP-flanked *Trp53* gene (*p53^LoxP/LoxP^*) or *p53^+/+^* and to a Rosa26::tdTom inducible reporter strain (Jax 007914) (Marino *et al*., 2000; Mori *et al*., 2006; Madisen *et al*., 2010). All animals were in a mixed 129xC57BL/6J background and both males and females were analysed between 7-10 weeks of age. Approximately an equal number of male and female mice were used per experiment.

### Neural progenitor cell culture

NPCs were isolated from the SVZ of postnatal day 9-12 (P9–P12) mouse brains as previously described (Ottone *et al*., 2014). Briefly, following microdissection, the SVZ was digested by incubation in HBSS (Invitrogen, 14170-088) supplemented with 0.25% trypsin and 60 U ml^−1^ Dnase I (Sigma, D4263) for 2 minutes at 37°C. Single cell suspensions were plated onto poly-L-lysine (PLL)-coated plates in SVZ explant medium consisting of DMEM/F12 (Invitrogen, 11320074), 3% FBS (Invitrogen), 20 ng ml^−1^ EGF (Peprotech, 315-09-1000) for 48 h. NPCs were routinely grown, for up to 8–10 passages, in SVZ culture medium consisting of DMEM/F12 (Life technologies, 11320074), 0.25% FBS, N2 (Life technologies, 17502001), 20 ng ml^−1^ EGF, 10 ng ml^−1^ bFGF (Peprotech, 450-33A) and 35 μg ml^−1^ bovine pituitary extract. For the *Cdkn1a* loss-of-function experiments, NPCs were isolated from the SVZs of *p21^+/+^* or *p21^-/-^* mice, a kind gift of Dr Owen J. Sansom (Cole *et al*., 2010).

### bmvEC culture

C57BL/6J primary mouse brain microvascular endothelial cells (bmvEC) were purchased from Cell Biologics (C57-6023) and subcultured on plates coated with attachment factor protein (Life Techonologies, S006110) in Endothelial Cell Growth Medium (Promocell, C-22111).

### Tamoxifen, EdU, Fish oil and Etomoxir administration

Tamoxifen (Sigma, T5648) was administered to 7-9 weeks-old mice by intraperitoneal injection (i.p.) at 100mg/kg/d. For assessment of resting type-B cells, EdU (Santa Cruz, sc-284628) was administered in the drinking water (0.2 mg/ml) *ad libitum* for 7 days, followed by tamoxifen administration for five consecutive days and mice were sacrificed 3 days later. To examine dormant type-B cells, EdU was administered in the drinking water *ad libitum* for 14 days. Mice were injected with tamoxifen over the last 5 days of EdU labelling and sacrificed 24h later. For Omega-3 FA administration experiments, 6-weeks old mice were given daily 100μL of fish oil (Sigma, F8020) corresponding to 0.2-0.43 g/kg/d docosahexaenoic acid (DHA) and 0.4-0.67 g/kg/d eicosapentaenoic acid (EPA) by oral gavage for 3 weeks. Tamoxifen was administered over the last 5 days of fish oil administration and SVZs collected 24h later. To prevent oxidation of the fish oil, aliquots were protected from direct light and supplemented with 40μM EDTA and 0.5mg/ml Ascorbyl Palmitate (Sigma, PHR1455). Control 6-weeks old mice received 100μL of PBS containing 40μM EDTA and 0.5mg/ml Ascorbyl Palmitate. For the tumour initiation experiments, we used a *de novo* somatic GBM model based on CRISPR/Cas9-mediated deletion of *Nf1, Pten* and *Trp53* tumour suppressors (NPP), as described (Brooks *et al*., 2021; Garcia-Diaz *et al*., 2021). Briefly, sgRNAs (single guide RNA) were expressed from a PiggyBac vector alongside a tdTomato reporter to fluorescently label resulting tumours. PiggyBase transposase was co-expressed with Cas9 in a second non-integrating plasmid. NPP-*Pparpgc1a-OE* and NPP-*Ppara*-KO constructs were generated using InFusion Kit (Clontech, 638917) and T4 DNA Ligase (NEB, M0202S), following manufacturer’s instructions. To generate the NPP-*Ppara*-KO, sgRNA to target *Ppara* (5’-GCCGGGGGACTCGTCCGTGC-3’), as reported (Sanjana, Shalem and Zhang, 2014), was cloned in the NPP plasmid 3’ of the *Nf1, Pten, Trp53* sgRNAs. Mouse *Ppargc1a* CDS was cloned after the tdTomato sequence as a polycistronic construct with a T2A linker for the generation of the NPP-*Pparpgc1a*-OE plasmid (Brooks *et al*., 2021; Garcia-Diaz *et al*., 2021). C57BL/6 female mice were randomly placed on an *ad libitum* fish oil supplemented diet (Teklad Global 2020X diet supplemented with 30-32g fish oil/kg containing 13.5% EPA and 10.5% DHA) or control diet (Teklad Global 2020X diet) following the observation of a copulation plug. Pups from these females were injected at P2 with the two plasmids mixed at a ratio of 1:1 using a Femtojet microinjector (Eppendorf) directly into the lateral ventricle. Plasmids were electroporated into the sub-ventricular zone using the Gemini X2 Generator set for 5 square pulses, 50 msec/pulse at 100 volts, with 950 msec intervals (Feliciano et al, 2013). Mice were maintained on fish oil supplemented diet or control diet until collection. After 3.5 weeks animals were injected with EdU (5mg/kg) 2 hours prior to sacrifice by transcardial perfusion of paraformaldehyde (PFA, 4%) under terminal anaesthesia. The brains were collected, stored overnight in PFA at 4°C prior to Vibratome sectioning (50μm) and subsequent analysis of tumour development. For the inhibition of FAO *in vivo*, pups were treated with etomoxir (Sigma-E1905) starting 10 days after electroporation and continuing for 2 weeks until sacrifice. This was achieved through lactation via etomoxir i.p. injections of dams at 10mg/kg every other day during the first week and through i.p. etomoxir injection of the pups at 5mg/kg every other day during the second week.

### FACS

qNSCs and aNSCs from the SVZ were FACS purified from GFAP::GFP mice as previously described (Codega *et al*., 2014) using the following antibodies: anti-mCD24 PE (1:2000; BD Pharmingen 12-0242-82), biotin conjugated anti-mCD133 (1:100, eBioscience 13-1331-82) and PE-Cy7 conjugated streptavidin (1:1000, eBioscience 25-4317-82) and EGF-complexed to Alexa Fluor™ 647 (1:100, ThermoFisher E35351). For isolation of *p53^wt^* and *p53^icKO^* qNSCs, mice were sacrificed 24h after tamoxifen administration. Upon dissociation, cells were stained with anti-mCD24-eFluor450 (1:2000; eBioscience 48-0242-82), biotin conjugated CD133 (1:100) and PE-Cy7 conjugated streptavidin (1:1000) and EGF-complexed to Alexa Fluor™ 647 (1:100). Zombie Green™ Fixable Viability Kit (1:1000, Biolegend 423111) or DAPI (1:10000, Sigma D9542) were used to assess cell viability. For both sets of experiments cell populations were manually gated based on FMOs, sorted using FACS Aria III (BD Biosciences) and collected into RLT buffer for RNA extraction.

### Co-culture experiments and cell treatments

Co-culture experiments were performed as previously described and analysed 48h later (Ottone *et al*., 2014). In brief, bmvEC were plated at a density of 2.6×10^4^cells/cm^2^ The following day, NPCs were plated either alone or onto endothelial monolayers at a density of 34×10^3^cells/cm^2^ After 48h, the NPCs were removed by selective trypsinization before analysis. Treatments were as follows: PPARα agonist WY-14643 was purchased from Sigma (C7081) and added to NPCs alone or in co-culture with bmvECs at the concentration of 200μM for 24h. For the EdU incorporation assay, 10μM EdU was added to the culture media for 2h prior to fixation and detection using the Click-iT™ EdU Alexa Fluor™ 647 Flow Cytometry Assay Kit (Thermo Fisher Scientific, C10424), according to manufacturer’s instructions. DNA was stained with DAPI. The cells were analysed on Fortessa X20s flow cytometer (BD). For *p53* loss-of-function experiments, *p53^loxP/loxP^* NPCs were infected with adenoviruses expressing a codon-improved Cre (iCre) under a CMV promoter (Ad-CMV-iCre, Vector Biolabs, #1045). Adenoviruses carrying CMV promoter only were used as controls (Ad-CMV-Null, Vector Biolabs, #1300). NPCs were used within 2-3 passages post infection to minimise compensatory effects of p53 loss. For luciferase assays, NPCs were nucleofected with a reporter plasmid containing the p53 binding element (PG13-CAT) prior to seeding in alone and bmvEC coculture for 48h (Kern *et al*., 1992; El-Deiry *et al*., 1993). Luminescence was normalized to viable cell numbers in each condition.

### Immunohistochemistry and Immunofluorescence

SVZ wholemounts were prepared as previously described (Mirzadeh *et al*., 2010). For Ascl-1 staining, mice were perfused with normal saline prior to dissection to reduce background from endogenous IgG in the blood. SVZs were fixed overnight in 4% PFA at 4°C, permeabilised for 1.5h in blocking solution, consisting of PBS with 10% donkey serum and 2% Triton X-100 at RT and incubated for 48h in primary and secondary antibodies diluted in PBS containing 2% TritonX-100 and 10% donkey serum at 4°C. Primary antibodies were: rabbit anti-Ki67 (1:50, abcam ab16667) and mouse anti-Ascl1 (1:200, a kind gift from Francois Guillemot (Urbán *et al*., 2016). EdU was detected using Click-iT® EdU Imaging Kit (ThermoFisher Scientific, C10340). For immunofluorescence experiments, cells were fixed in 4% PFA, permeabilised in 0.5% Triton X-100, blocked in PBS containing 10% serum for 1h and incubated in primary antibody diluted in blocking buffer O/N at 4°C. Primary antibodies were: mouse anti-p53 (1:500, Cell Signaling 2524), rabbit anti-GFAP (1:1000, Dako Z0344), mouse anti-SOX2 (1:100, abcam ab79351), mouse anti-NESTIN (1:200, santa cruz sc-33677), mouse anti-PPARα (1:25, Arigo Biolabs ARG55240). Alexa Fluor conjugated secondary antibodies (ThermoFisher Scientific) were diluted in 10% serum in DAPI (1:10000 in PBS) and incubated at RT for 1h. CellTracker™ Green (ThermoFisher Scientific, C7025) was used to label NPCs prior to seeding. Imaging was carried out using the Zeiss LSM880 confocal microscope. Quantifications were performed by using Fiji ImageJ (Schindelin *et al*., 2012). For all wholemount quantifications, percentages of type-B and type-C were calculated over the total number of tdTom^+^ type-B and type-C cells.

### Quantitative RT-PCR

For in vitro experiments, RNA was extracted using RNeasy mini kit (Qiagen, 74104) following the manufacturer’s instructions. RNA was reverse transcribed using iScript gDNA clear cDNA synthesis kit (Bio-rad, 1725034) and quantitative PCR was performed using the qPCRBIO SyGreen Mix Lo-Rox (PCR Biosystems, PB20.11). For assessment of acutely FACS-purified qNSCs and aNSCs, RNA was extracted using RNeasy Plus MicroKit (Qiagen, 74034) according to the manufacturer’s instructions and cDNA libraries were prepared using the Smart-seq2 protocol (Picelli *et al*., 2013). Primers used were:

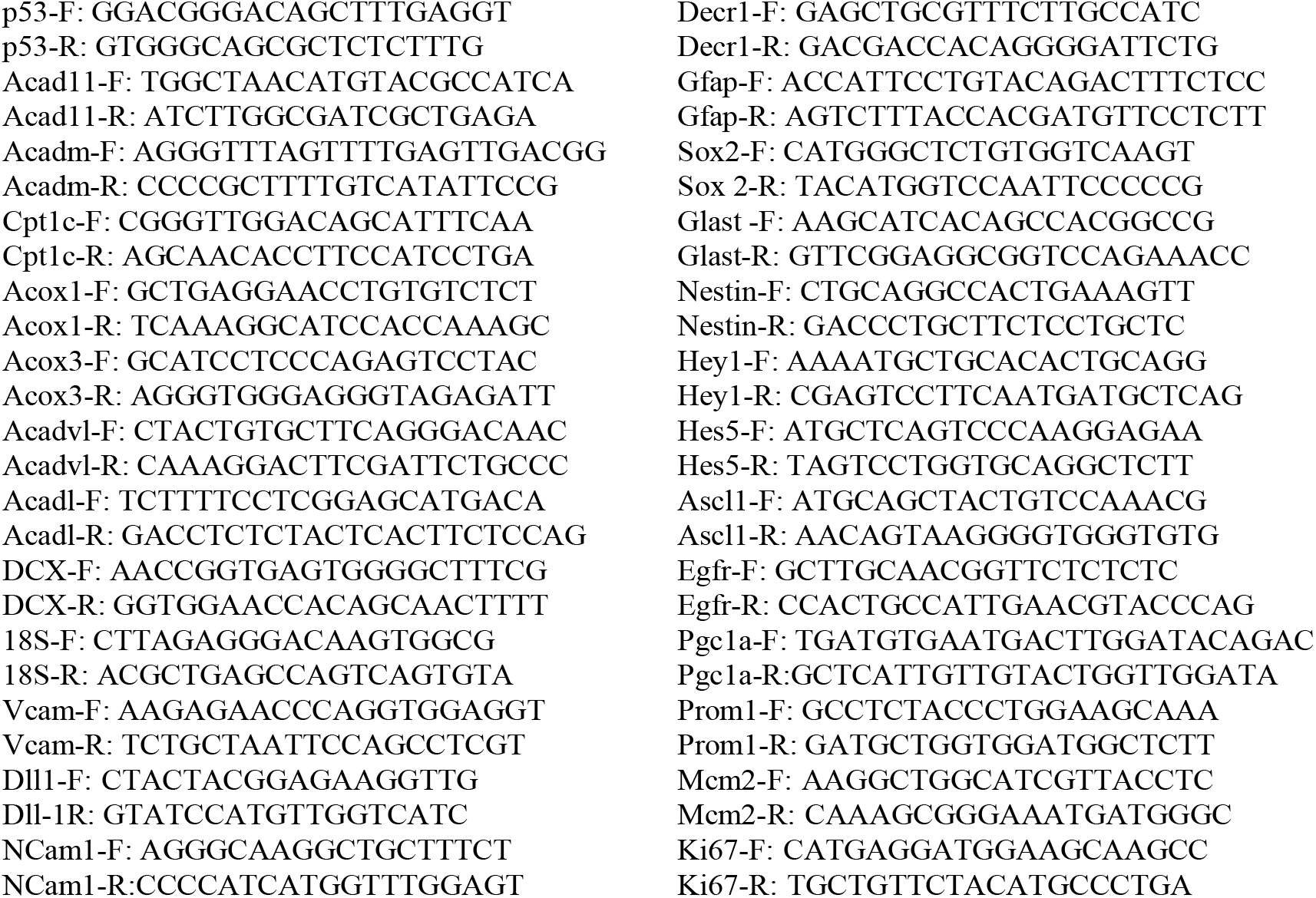

### FAO assay

Fatty acid oxidation was measured using the commercially available Fatty Acid Oxidation (Abcam, ab217602) and Extracellular Oxygen Consumption Assay kits (Abcam, ab197243) according to the manufacturer’s instructions. Treatment with 25μM FCCP served as positive and samples without cells as negative controls, respectively. Extracellular O2 consumption was measured every 90 seconds for 2h at Ex/Em=380/650 nm using the VarioSkan LUX Plate reader set at 37°C. (ThermoFisher Scientific). Oxygen Consumption Rate (OCR) was determined by calculating the slope of the linear proportion of the signal profiles.

### ^13^C-palmitate tracer experiments and GC-MS analysis

A 100mM stock of [U-^13^C]-palmitic acid (Sigma, 605573) was prepared in ethanol. This was diluted 100-fold into DMEM/F12 media containing 2% w/v fatty acid-free BSA (Sigma A8806) and sonicated for 15 minutes in a water bath sonicator at room temperature to form soluble BSA/palmitate complexes. The resulting media contained 1mM [U-^13^C]-palmitic acid, 2% BSA and 1% ethanol (170 mM) with a palmitate:BSA molar ratio of 3.3:1. Following filtration through a 0.2 μm PES filter (Millipore, Burlington, MA) this was diluted 1:10 in SVZ culture medium, giving a final concentration of 100μM [U-^13^C]-palmitic acid.

Cells in co-culture, in the presence or absence of PPARα agonist WY-14643 (200μM), were incubated with 100μM U-^13^C-palmitate SVZ medium for 6 hr or 24 hr. Cell were selectively dissociated from the endothelial monolayer and rapidly quenched on dry ice/ethanol. Samples were then pelleted at 0°C, washed with ice-cold PBS pH 7.4, transferred to 2 mL tubes and metabolites extracted at 4°C for 1 hr (with 3 × 8 min sonications in a water bath sonicator) using 600 mL chloroform/methanol (2:1, v/v, containing ^13^C lauric acid internal standard at 5nmol per sample). Extracts were transferred to 1.5 mL tubes and dried in a SpeedVac. The pellet was then re-extracted with 450 mL methanol/water (2:1, v/v, containing 1 nmol scyllo-Inositol internal standard; 4°C, 8 min sonication), the extract was combined with the first extract and then re-dried. Polar and apolar metabolites were separated by phase partitioning with chloroform/methanol/water (1:3:3, v/v/v) and then analyzed by GC-MS.

GC-MS data acquisition was performed largely as previously described (MacRae *et al*., 2013), using an Agilent 7890B-5977A GC-MSD in EI mode after derivatization of twice methanol-washed dried extracts by the addition (a) for polar metabolites of 20 μL of a 20 mg/mL solution of methoxyamine hydrochloride in pyridine (both Sigma) at room temperature for >16 hr and 20 μL BSTFA + 1% TMCS (Sigma) at RT for >1 hr, or (b) for fatty acids of 25 μL chloroform/methanol (2:1, v/v) and 5 μL MethPrepII (Grace Alltech) at room temperature with no incubation. GC-MS parameters were as follows: carrier gas, helium; flow rate, 0.9 mL/min; column, DB-5MS (Agilent); for polar analyses: inlet, 270°C; temperature gradient, 70°C (2 min), ramp to 295°C (12.5°C/min), ramp to 320°C (25°C/min, 3 min hold); for apolar analyses: inlet, 250°C; temperature gradient, 70°C (1 min), ramp to 230°C (15°C/min, 2 min hold), ramp to 325°C (25°C/min, 3 min hold). Scan range was m/z 50-550 (polar) and 50-565 (apolar). Data were acquired using MassHunter software (version B.07.02.1938). Data analysis was performed using MANIC software, an in house-developed adaptation of the GAVIN package (Behrends, Tredwell and Bundy, 2011). Metabolites were identified and quantified by comparison to authentic standards, and label incorporation estimated as the percentage of the metabolite pool containing one or more ^13^C atoms after correction for natural abundance.

### RNA-Sequencing

For RNA sequencing RNA, was isolated using RNeasy mini kit (Qiagen, 74104) according to manufacturer’s instructions. Libraries were prepared using the Truseq mRNA stranded kit and quality control checks were performed by Qubit and Bioanalyser analysis. Libraries were then pooled and run on a MiSeq Nano Flow Cell (V2 reagents) (Single Read 26 cycles) to check the balance of the libraries within the pool and balance adjusted where necessary. Clonal clusters of each library were then amplified onto an Illumina Flow Cell using the Illumina cBot system and sequenced on a HiSeq 2500 (v4 chemistry) as a Paired-End 100bp.

### RNA-seq data pre-processing and differential expression analysis

Raw reads were aligned to the mouse genome (NCBI Build 37, USCS mm9) using the TopHat v.2.0.11 software(Kim *et al*., 2013) and assigned to genomic features using HTSeq v.0.6.1(Anders, Pyl and Huber, 2015). Differential expression analysis was performed and normalized counts were generated using the DESeq2 Bioconductor package (Love, Huber and Anders, 2014) (Supplemental Table S1). RNA-seq data that support the findings of this study has been deposited in GEO (accession codes pending).

### CUT&RUN library preparation and sequencing

CUT&RUN experiments were performed as previously described (Skene, Henikoff and Henikoff, 2018). In brief, 5×10^5^ NPCs grown either in alone or in coculture conditions were attached to concanavalin A–coated magnetic beads (Bangs Laboratories, BP531) and incubated overnight at 4°C in 0.05% Digitonin-Antibody buffer containing a rabbit anti mouse-p53 Ab (1:100 Leica, NCL-p53-CM5p). Samples incubated with rabbit IgG (1:150) were used as control. After incubation, samples were washed in Dig-wash and incubated with ProteinA-MNase fusion protein at 700 ng/mL (EpiCypher, 15-1016-EPC) on a tube rotator at 4 °C for 1 hr. Chromatin was digested in Incubation Buffer containing 1M CaCl_2_ at 0 °C for 10 minutes, chromatin fragments were released by incubation at 37 °C for 30 min. Chromatin was purified by performing Phenol Chloroform Extraction. CUT&RUN barcoded libraries for Illumina sequencing were prepared using the NEBNext® Ultra™ II DNA Library Prep Kit for Illumina® (NEB, E7645) accordingly with Zhu et al.(Zhu *et al*., 2019). PCR amplification products were cleaned and size selected by using AMPure XP beads (Beckman Coulter, A63881). The resulting purified libraries were quantified by Qubit. Library size distribution was checked by using Agilent Bioanalyzer® High Sensitivity DNA chip (Agilent Technologies, 5067-4626). Indexed libraries were pooled and Illumina Paired-End (42×42bp, 6-bp index) sequencing was performed using Nextseq 500 platform with NextSeq 500/550 High Output Kit v2 (75 cycles).

### CUT&RUN data pre-processing and peak calling

Sequencing data was analysed using the CUT&RUNTools pipeline (Zhu *et al*., 2019). Briefly, Fastq files were trimmed using Trimmomatic (Bolger, Lohse and Usadel, 2014) before alignment to Mus musculus GRCm38 (mm10) genome with Bowtie2 (using the --dovetail option) (Langmead and Salzberg, 2012). Aligned reads were converted to bed format using bedtools bamtobed with -bedpe option, before filtering for fragments shorter than 120bp. Normalisation of each sample to mm10 read depth was performed using bedtools genomecov with a scale factor generated by division of an arbitrary large number by read depth for each sample. Peaks were called using MACS2 (Zhang *et al*., 2008). Downstream peak analysis was performed in R. Peaks were annotated using the Chippeakanno R package (Zhu *et al*., 2010) and visualised in IGV.

### Bioinformatics analysis

RNA-seq and CUT&RUN functional analysis was performed using custom R scripts. Hierarchical clustering was performed using the “pheatmap” package. Figure 2C: Genes with an absolute DEseq2 log_2_ ratio > 1 in one or more contrast and defined as aNSCs or qNSCs markers in one the three published studies were used (Codega *et al*., 2014; Llorens-Bobadilla *et al*., 2015; Mizrak *et al*., 2019). Figure 2D: GO term enrichment analysis of Figure 2C clusters was performed using VLAD (http://proto.informatics.jax.org/prototypes/vlad/) (Richardson and Bult, 2015) Figure 2E: Genes with an absolute DEseq2 log_2_ ratio > 0.5 and adjusted p-value < 0.5 in the KO AL/WT AL or KO CO/WT CO contrasts and belonging to the fatty acid metabolic process GO category (GO_0006631) were used. Figure S2A: Genes with an absolute DEseq2 log_2_ ratio > 1 in the WT CO/WT AL contrast were used. Figure S2B-C: Genes with an absolute DEseq2 log_2_ ratio > 1 (up) or < −1 (down) and adjusted p-value < 0.05 in the WT CO/WT AL contrast were used for functional analysis using the g:Profiler package (Figure S2B, https://biit.cs.ut.ee/gprofiler/gost) (Raudvere *et al*., 2019), or VLAD ( Figure S2C, http://proto.informatics.jax.org/prototypes/vlad/) (Richardson and Bult, 2015). Figure 3A: Genes significantly regulated (DESeq2 adjusted p-value < 0.05) in the WT CO/WT AL contrast and found to be targets of *Ppara* but not *Ppard* or *Pparg*, or targets of *Ppard* but not *Ppara* or *Pparg* were used (Fang *et al*., 2016). Figure S3A: Genes significantly regulated (DESeq2 adjusted p-value < 0.05) in either the KO AL/WT AL or KO CO/WT CO contrasts and showing significant p53 binding in at least one of four CUT&RUN experiments (MACS Q value < 0.01) were used (Zhang *et al*., 2008).

### Quantification and statistical analysis

Statistical analysis was performed using GraphPad7 or GraphPad9 built in tools. All graphs represent the mean ±SEM. Significance is stated as follows: p>0.05 (ns), p<0.05 (*), p<0.01 (**), p<0.001 (***), p<0.0001 (****). Two-tailed Student’s t test was used for statistical comparisons between two groups. Two-way ANOVA with Tukey’s multiple comparisons was used to determine statistical significance of multiple comparisons. Statistical details of experiments can be found in the figure legends.

