## Supplemental file for "Diet suppresses tumour initiation by maintaining quiescence of mutation-bearing neural stem cells"

Amodeo et al., Supplemental Figure 1

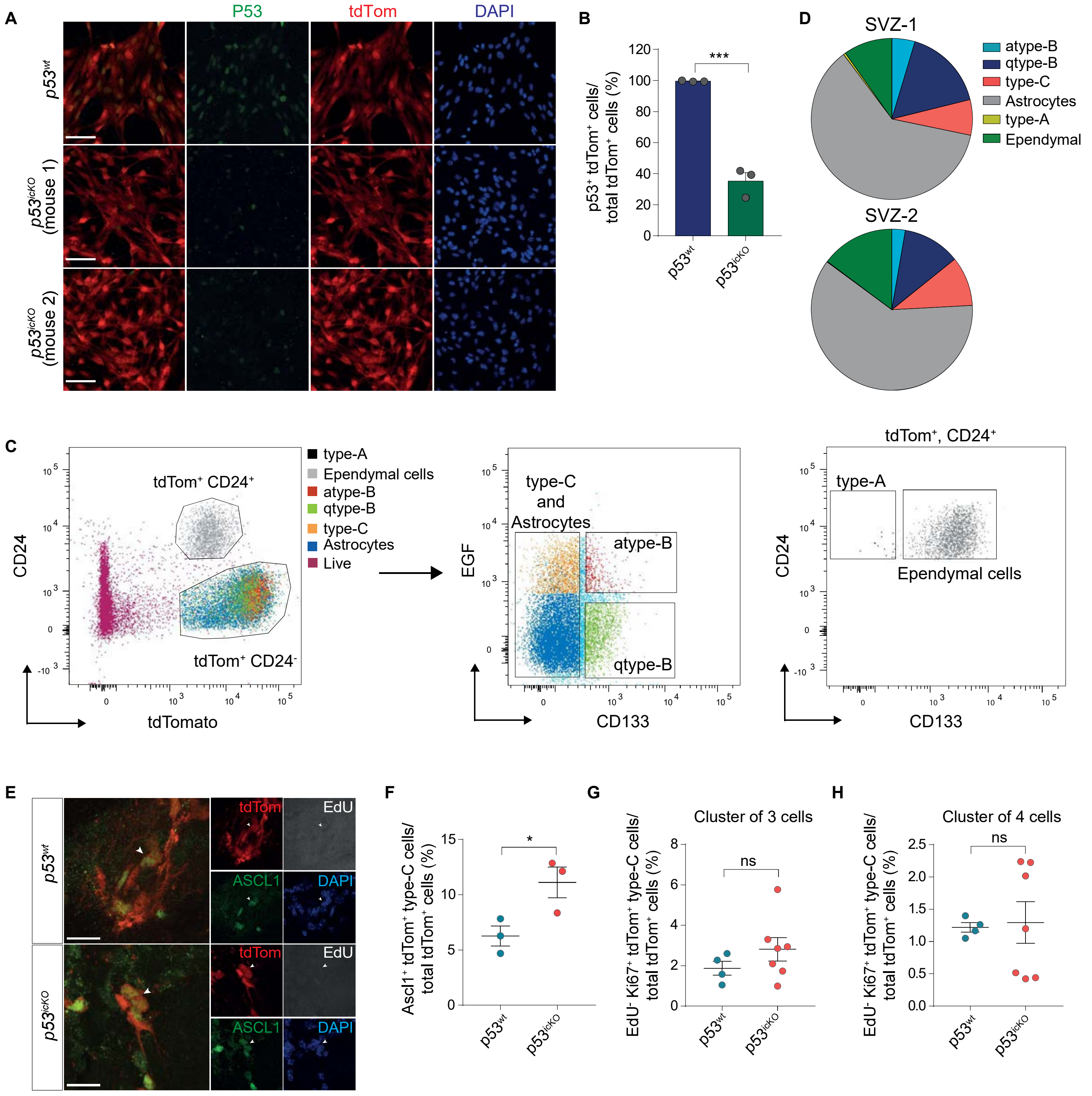

### Supplemental Figure legends

**Supplemental Figure 1 (Related to Figure 1). Selectivity of the *$p53^{icKO}$* model. A,** Representative fluorescence images showing p53 expression in NPCs isolated from the SVZ of adult mouse brain 24h after tamoxifen administration. p53 is shown in green, recombined NPCs are $tdTom^+$ and nuclei are counterstained with DAPI (blue). Scale bar=100µm. **B,** Quantification of the percentage of $p53^+$ NPCs over the total number $tdTom^+$ NPCs. Mean±SEM, $p53^{wt}$ n=3, $p53^{icKO}$ n=3. ***p<0.001, unpaired two-tailed Student t-test. **C,** Representative FACS plots showing the gating strategy for all $tdTom^+$ SVZ subpopulations in $p53^{icKO}$ mice. $tdTom^+/CD24^-$ cells were selected and gated on EGF-AF647 and CD133-PE-Cy7 levels to define $tdTom^+/CD24^-/EGFR^+/CD133^+$ aNSC cells, $tdTom^+/CD24^-/EGFR^-/CD133^+$ qNSCs and $tdTom^+/CD24^-/EGFR^-/CD133^-$ and $tdTom^+/CD24^-/EGFR^+/CD133^-$ astrocytes. Proportion of type-C cells was estimated from the $tdTom^+/CD24^-/EGFR^{high}/CD133^-$ population and distinguished from astrocytes based on cell size. Ependymal cells were defined from the $tdTom^+/CD24^+$ population as $tdTom^+/CD24^+/CD133^+$ cells and type-A as $tdTom^+/CD24^+/CD133^-$ cells. Cell viability was assessed with Viability Dye Zombie Green. Single immuno-stained and Fluorescence Minus One (FMO) SVZ cell suspensions were used as controls. $tdTom^-$ cell suspension served as negative control. **D,** Pie chart showing percentages of each subpopulation in $p53^{wt}$ 24h post-recombination. **E,** Representative fluorescence images of $Ascl1^+/tdTom^+$ type-C cells in SVZ wholemounts of $p53^{wt}$ and $p53^{icKO}$ mouse brains 3 days post-recombination. White arrowheads denote pairs of $Ascl1^+/tdTom^+$ type-C cells. Scale bar=20µm. **F,** Quantification of the percentage of $Ascl1^+/tdTom^+$ type-C cell pairs over the total number of $tdTom^+$ type-B and type-C cells. Mean±SEM, $p53^{wt}$ n=3, $p53^{icKO}$ n=3. *p<0.05, unpaired two-tailed Student t-test. **G-H,** Quantification of the percentage of

EdU$^-$/Ki67$^+$/tdTom$^+$ clusters of 3 type-C cells (G) and 4 type-C cells (H) in SVZ wholemounts from $p53^{wt}$ and $p53^{icKO}$ mice subjected to experimental protocol described in Fig.1A for the detection of resting qNSCs. Mean±SEM, $p53^{wt}$ n=4, $p53^{icKO}$ n=7. ns=not significant, unpaired two-tailed Student t-test.

Amodeo et al., Supplemental Figure 2

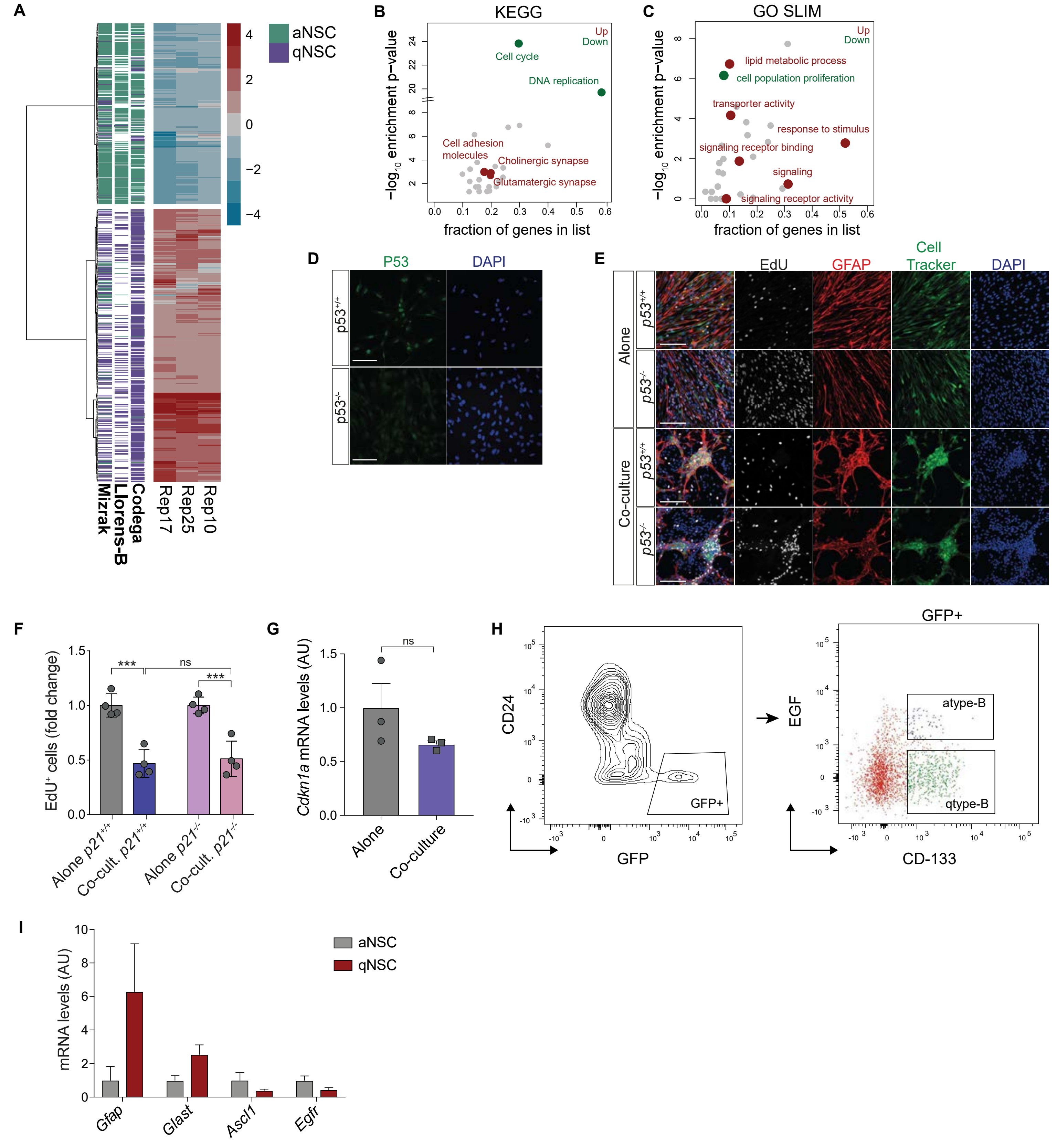

**Supplemental Figure 2 (Related to Figure 2). p53-mediated cell cycle arrest is p21-independent. A,** Hierarchical clustering of RNA-seq $\log_2$ expression ratios for three independent NPC co-culture experiments relative to alone controls shown alongside published data-sets from type-B cells FACS acutely purified from the V-SVZ (Codega *et al.*, 2014; Llorens-Bobadilla *et al.*, 2015; Mizrak *et al.*, 2019). **B-C,** KEGG and GO pathway analysis of genes up- and down-regulated upon co-culture. Enrichment p-values are shown as a function of the fraction of regulated genes present in KEGG pathways (B) and GO categories (C). Selected terms enriched in genes up- (red) and down-regulated (green) upon co-culture are highlighted. **D,** Representative immunofluorescence p53 staining (green) in NPCs infected with an adenovirus expressing Cre recombinase ($p53^{-/-}$) or a control adenovirus ($p53^{+/+}$). DAPI-stained nuclei are in blue. Scale bar=100µm. **E,** Representative EdU staining of $p53^{+/+}$ and $p53^{-/-}$ NPCs cultured alone or with bmvECs for 48h and pulsed with EdU for 2h prior to collection. NPCs are cell-tracker labelled (green) and stained for GFAP (red), EdU (grey) and DAPI (blue). Scale bar=20µm. **F,** EdU FACS profiles of $p21^{+/+}$ and $p21^{-/-}$ NPCs cultured alone or with endothelial cells for 48h and pulsed with EdU for 2h. Fold changes normalized to the respective alone controls are shown for each genotype. Mean±SEM, n=4 independent experiments. ns=not significant, ∗∗∗$p<0.001$, Two-way ANOVA with Tukey's multiple comparisons test. **G,** Quantitative RT-PCR analysis of *Cdkn1a* mRNA levels in NPCs alone and in co-culture. Fold changes relative to alone control are shown. Mean±SEM, n = 3, ns=not significant, unpaired two-tailed Student t-test. **H**, Representative FACS plots of the strategy used to prospectively purify SVZ subpopulations using GFAP::GFP mice as in Codega et al (Codega *et al.*, 2014). Cell viability was assessed with DAPI. Single immuno-stained SVZ cell suspensions from wild-type mice were used to set FACS gates. **I,** Quantitative RT-qPCR analysis of the mRNA levels of *Gfap*, *Glast*, *Ascl1* and *Egfr* in qNSCs and aNSCs cells FACS-purified as in H confirms successful enrichment of each subpopulation. Data represent the average of 3 independent sorts; Mean±SEM.

Amodeo et al., Supplemental Figure 3

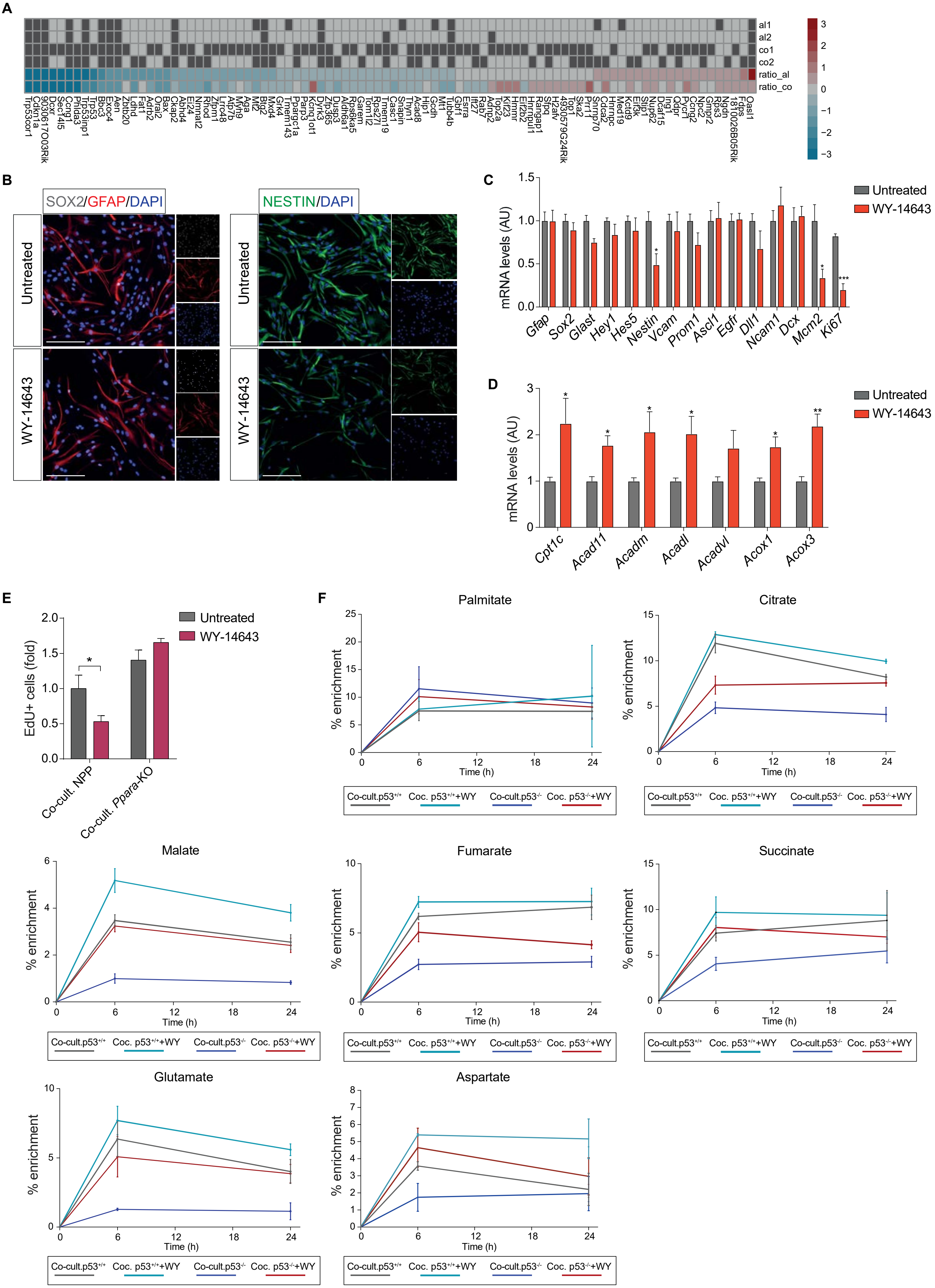

**Supplemental Figure 3 (Related to Figure 3). Validation of CUT&RUN experiments and WY-14643 treatment. A,** List of genes regulated upon p53 deletion in alone or co-culture conditions, which show significant binding by p53 in CUT&RUN analysis. Rows 1-4: alone (al) and co-culture (co) CUT&RUN experiments for two independent replicates are shown and dark grey rectangle represents significant p53 binding. Rows 5-6: RNA-seq $\log_2$ expression ratios between and *p53*$^{-/-}$ and *p53*$^{+/+}$ NPCs in alone (ratio_al) and in co-culture (ratio_co). **B,** Representative fluorescence images of SOX2 (grey), GFAP (red) and NESTIN (green) in NPCs untreated or treated with WY-14643 for 24h. Nuclei are counterstained with DAPI (blue). Scale bar=100µm. Note that PPARα agonist treatment does not induce NPC differentiation. **C-D,** Quantitative RT–PCR analysis of markers for qNSC (*Gfap*, *Sox2*, *Glast*, *Hey1*, *Hes5*, *Nestin*, *Vcam*, *Prom1*), aNSC/TAPs (*Ascl1*, *Egfr*, *Dll1*, *Ncam1*, *Dcx*), proliferation (*Mcm2 and Ki67*) and FAO genes (*Cpt1c*, *Acad11*, *Acadm*, *Acadl*, *Acadvl*, *Acox1* and *Acox3*) (D) in NPCs after 24h of WY-14643 treatment. Data are shown as fold change relative to untreated NPCs (control). Mean±SEM, n=5 independent experiments, *p<0.05, **p<0.01, ***p<0.001, unpaired two-tailed Student t-test. **E**, Quantification of EdU incorporation in NPP and NPP-*Ppara*-KO NPCs co-cultured with bmvEC for 24 h in the presence or absence of WY-14643 and pulsed with EdU for 2h prior to collection. Mean±SEM, n=3 replicates, *p<0.05, Two-way ANOVA with Sidak's multiple comparisons test. **F**, $^{13}$C enrichment following 6h or 24h labelling with [U-$^{13}$C]-palmitic acid. Near isotopic steady-state enrichments are observed for palmitate, tricarboxylic acid (TCA) cycle intermediates and the amino acids glutamate and aspartate. *p53*$^{+/+}$ and *p53*$^{-/-}$ NPCs in co-culture untreated and treated with the PPARα agonist WY-14643 for 24h were used. Graphs shows the % of $^{13}$C enrichment for all isotopologues of each metabolite at 6h and 24h.

Amodeo et al., Supplemental Figure 4

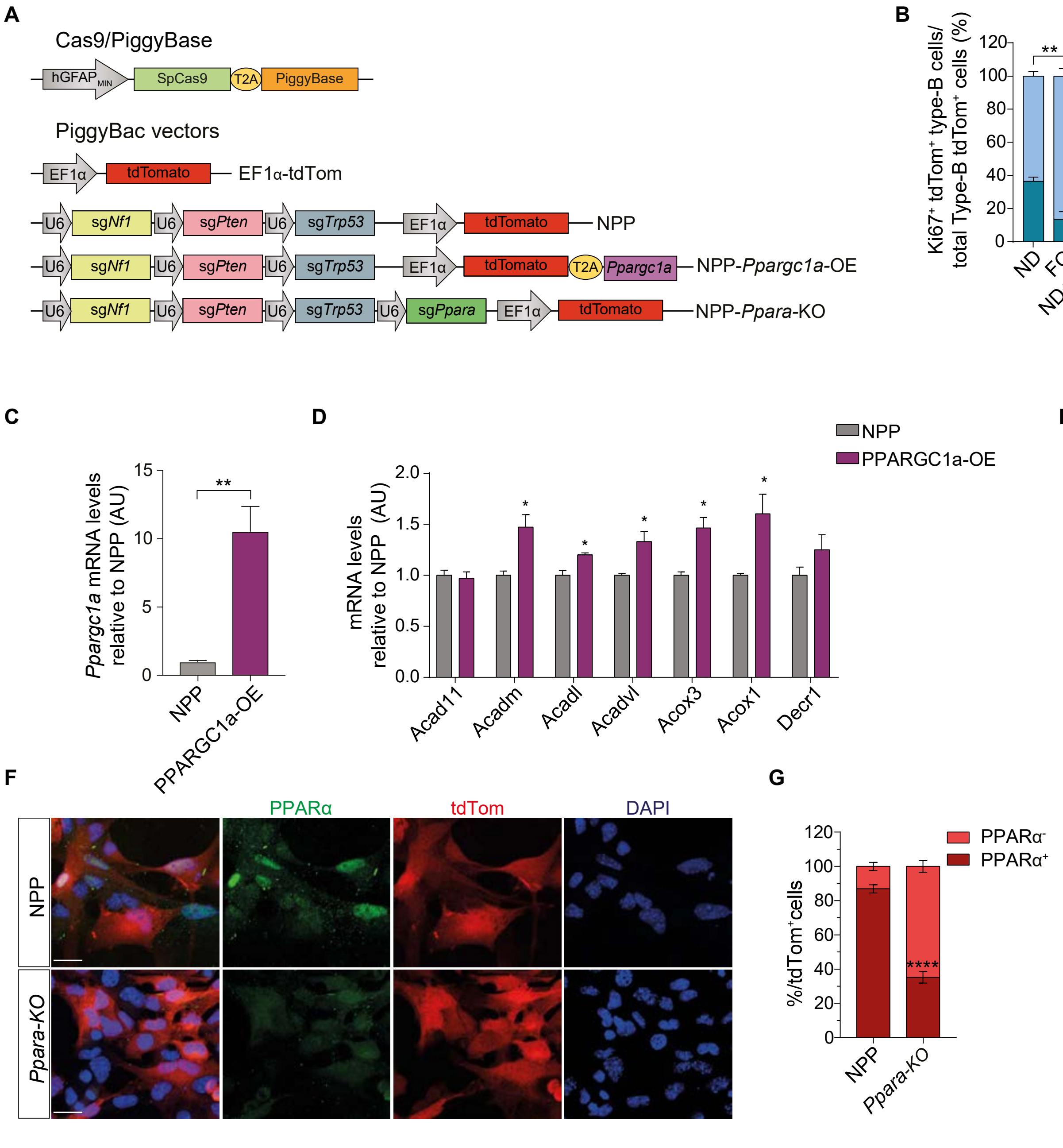

**Supplemental Figure 4 (Related to Figure 4). Validation of PPARGC1a overexpression and PPARα genetic deletion. A**, Schematic of the Cas9/PiggyBase and PiggyBac plasmids used to generate *Nf1, Pten and Trp53* knock-out tumours (NPP), or NPP tumours overexpressing *Ppargc1a* (NPP-*Ppargc1a*-OE) or knock-out for PPARα (NPP-*Ppara*-KO). **B,** Quantification of quiescent (Ki67$^-$/tdTom$^+$) and active (Ki67$^+$/tdTom$^+$) type-B cells in the SVZ of NPP tumour-bearing mice fed a normal or fish oil-supplemented diet in the presence or absence of etomoxir (ETO). Mean±SEM, **p<0.01, ***p<0.001, ****p<0.0001, Two-way ANOVA with Sidak's multiple comparisons test. **C-D**, Validation of *Ppargc1a* overexpression by qRT-PCR analysis (C) and analysis of FAO genes expression (D) in NPCs transfected with the NPP-*Ppargc1a*-OE plasmid. NPP transfected cells were used as control. Fold changes relative to NPP cells are shown. Mean±SEM, n=3 independent experiments, *p<0.05, **p<0.01, unpaired two-tailed Student t-test. Note that the majority of FAO genes are induced by PPARGC1a overexpression. **E**, Quantitative RT-PCR analysis to assess *Ppargc1a* overexpression in tdTom$^+$ FACS sorted cells isolated from tumours. Mean±SEM, n=2 independent experiments. **F**, Validation of PPARα deletion by immunocytochemistry. Shown are confocal images of tdTom$^+$ NPP and NPP-*Ppara*-KO cells isolated from tumours. PPARα is shown in green, transformed NPCs are tdTom$^+$ (red) and nuclei are counterstained with DAPI (blue). Scale bar=20µm. **G**, Quantification of the PPARα$^+$ and PPARα$^-$ cells from the fluorescence images showed in E. Data are shown as percentage of the total number of tdTom$^+$ cells. Mean±SEM, ****p<0.0001, unpaired two-tailed Student t-test.
